# How Hippocampus Solves Inverse Problems: A Circuit Hypothesis

**DOI:** 10.1101/2022.03.15.484469

**Authors:** David W. Arathorn

## Abstract

This paper proposes an explicit computational process which reproduces what are considered to be two essential experimentally observed hippocampus capabilities. These are path planning and determining the chain of ordered relationships which bind two concepts. These are both inverse problems. The paper presents algorithmic simulations of the process and a neuronal circuit (and variants) with the same dataflow, both consistent with the known neuroanatomy of the hippocampus. The process described here is purpose-built for hippocampus

## Introduction

The volume of research on hippocampus has revealed such a wide span of functions that a compact hypothesis of hippocampal function would seem nearly impossible. However, regularities in that functional span have been apparent for many years. Buzsaki observes that the signal dynamics across the hippocampal-entorhinal system also imply a common computation on a range of neocortical sources. [1] This paper proposes an explicit computational process which reproduces what are considered to be two essential experimentally observed hippocampus capabilities. These are path planning and determining the chain of relationships which bind two concepts. These are both inverse problems, not solvable by any of the varieties of neural nets. This paper presents algorithmic simulations of the process and a neuronal circuit with the same dataflow, both consistent with the known neuroanatomy of the hippocampus. The process described here is purpose-built for hippocampus function and architecture and does not derive from any pre-existing class of models.

Several key observations from the experimental literature underlie the hypothesis presented here. First, place cells in the hippocampus (Hc) can take on both spatial and entity definitions depending on their inputs from entorhinal cortex (EC), and the computations performed on the resulting representations in the Hc space are the same or very similar independent of those definitions. [2] Second, the primary computation performed by Hc is the establishment of a sequence connecting sets of place cells. [1] Third, such sequences can be captured in a memory structure that can be played back in the order of capture or the reverse of that order. [3] Fourth, the neuronal circuit that implements these computations is a loop comprised of two or three primary stages, and the computation by the circuit can involve multiple passes through the loop.

While the Hc literature spans many functions, two stand out as templates of much of the span. One is the navigational or route-planning function. [4] Another is the relation-establishing function, as in the classic transitive relation inferencing role. [5,6] Many of the other known or suspected functions of Hc are analogs of one or the other of these two, can use either the spatial or entity definitions of place cells, and can be addressed by the same spatial or relational computations. The somewhat different mechanics between the spatial and relational modes of operations, while operating within the same circuit, will be addressed explicitly in this paper.

Since the same structure is recruited for both navigational and relational computations, it is logical to assume one of these functions evolved first and the second was a modification of the first. It is hard to construct an argument against the notion that navigation came first. In any event navigation has simpler requirements and is easier to explain, so that mode will be presented first. Explanation of the relational mode will follow. These presentations will, for clarity and compactness, be couched in algorithmic terms first and then translated to a medium-level presentation of a circuit which can execute the algorithmic steps and is consistent with Hc neuro-architecture. And with that specific model and behavior well defined, reconciliation to biological evidence will be discussed.

### Navigation: Animation of Principle of Operation

The mechanics of navigation is best illustrated by frame-by-frame animations produced by a simple Matlab program. The principle is presented first in the context of a “toy” path-planning problem with starting and ending locations placed very close together and no intervening obstacles so that a few steps complete the path. This allows the computational process to be narrated compactly. Afterward, using the same program the animations are expanded to more a larger field of simulated place cells and realistic length paths, some including obstacle or stricter path constraints. The dataflow of the simulation is represented in Figure 1.

**Figure 1:**
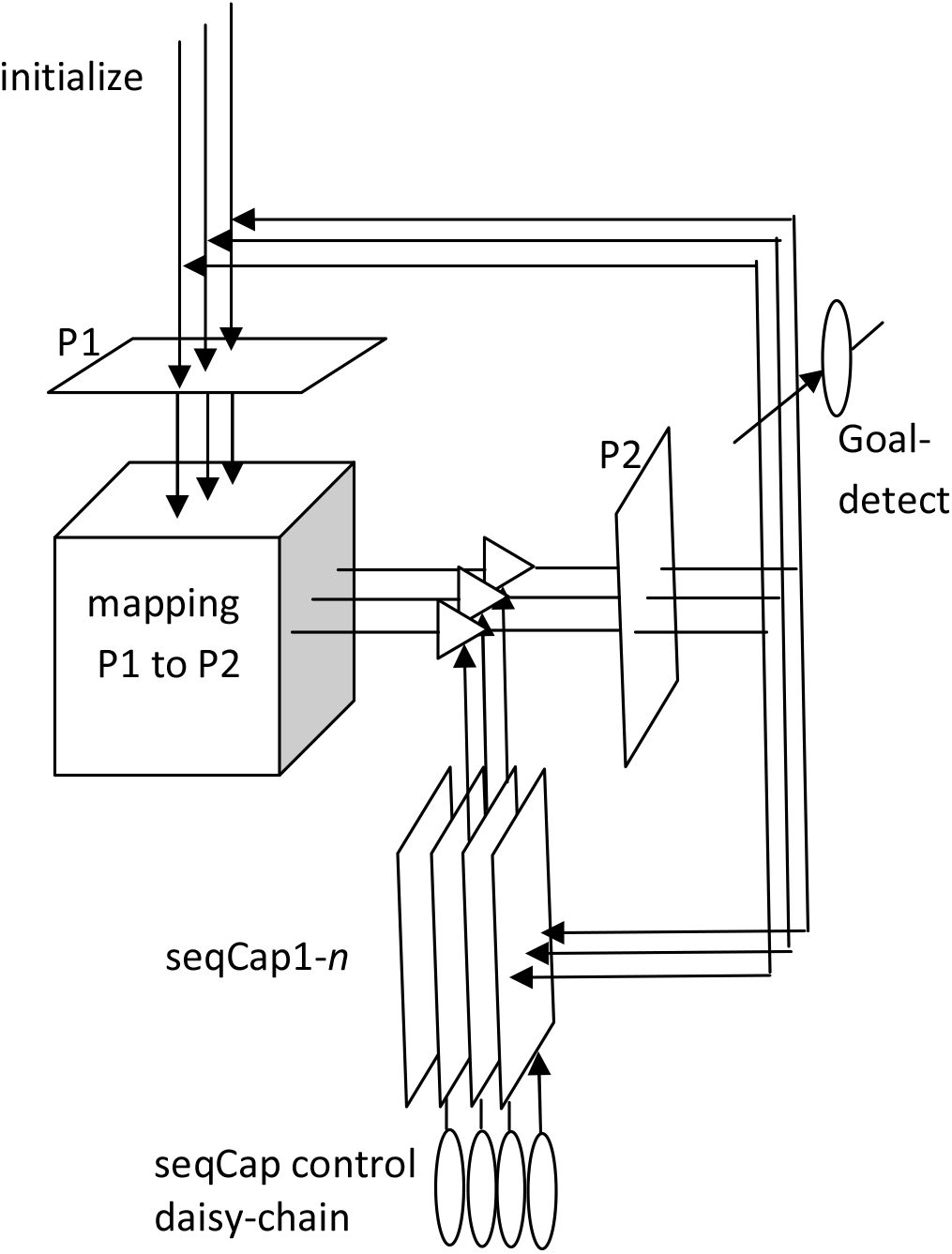
Dataflow of the simulation

Figure 1 shows 5 categories of cells. **P1** and **P2** are 2D place cell arrays. The connectivity between the axons of P1 and the dendrites of P2 implement a local mapping, to be described shortly. A set of *n* short-term memory or SequenceCapture arrays, **seqCap1-n**, provide inhibitory gating to the dendrites of P2. The axons of P2 loop back to the dendrites of P1 and also provide input to the seqCap1-n arrays. A set of n control cells, the **seqCap daisy-chain**, select in sequence which of the n seqCap arrays is providing and storing activation patterns to and from P2. The **goal-detect cell**, shown as single but likely several, monitors the axons of P2 and is triggered when a path goal is reached. The dendrites of P1 also provide an input for initialization of P1 from areas external to the circuit shown, e.g. EC.

The function and more detailed organization of the cells listed above are discussed in more detail below. Where there are alternative circuit architectures with the same function, that is also discussed. In the description of the toy problem simulation dynamics, the emphasis will be on the flow of activation that results in the path computation. Aspects of the control signals which sequence this computation will be mentioned but dealt with in more detail in the discussion of the circuits, much later in the paper.

The panels of Figure 2(a-b), and later Figure 2(c-f), render the activity of the principle simulated cell arrays for the toy problem. Each square represents a 60×60 field of place cells with the relative activity of each cell indicated by the color of the pixel representing the cell. Three arrays are displayed in each column representing the state of the P1, P2 and SeqCap arrays in Figure 1 at different time steps.

**Figure 2(a):**
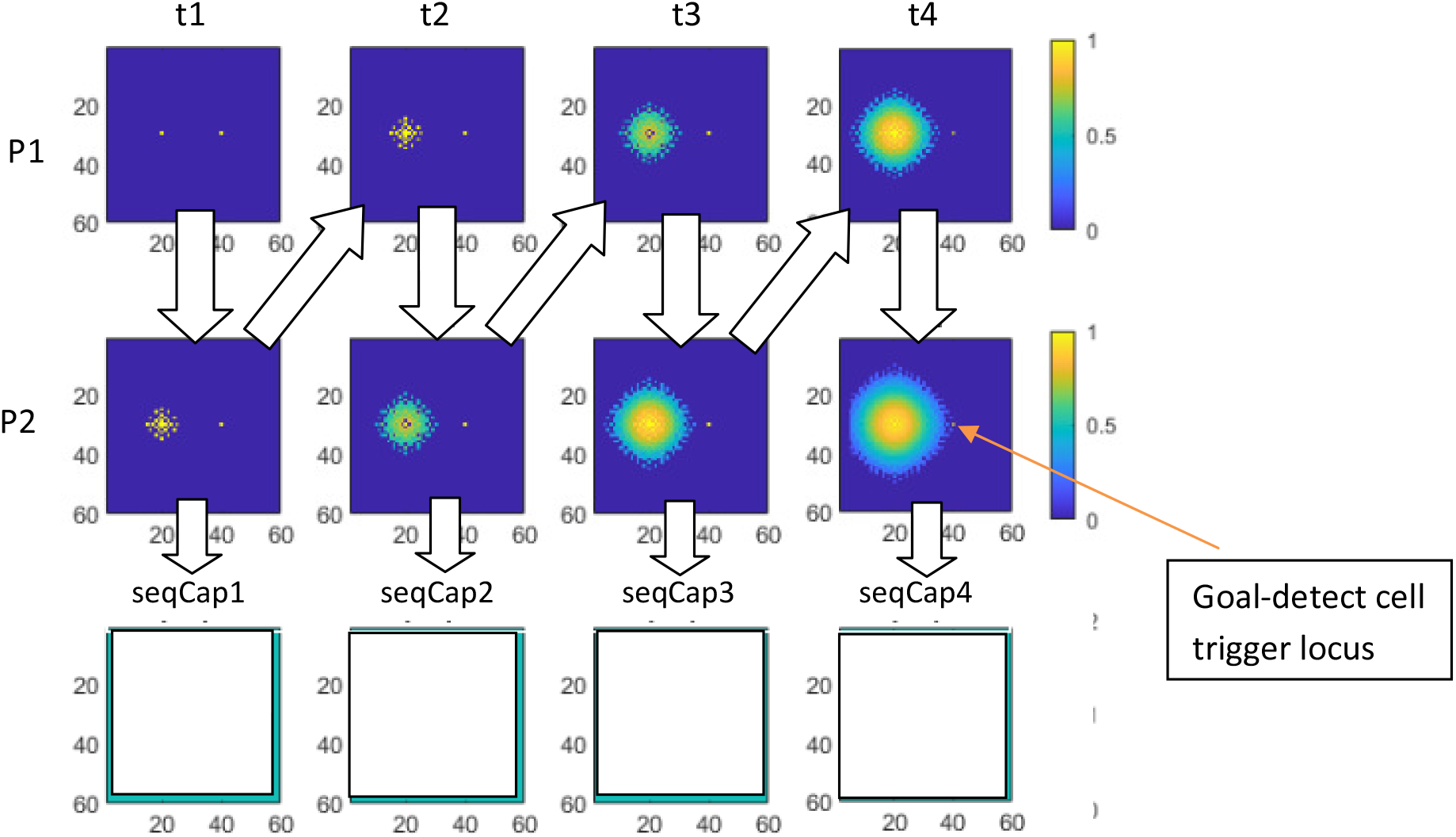
1^st^ pass forward (all figures compressed amplitude). seqCap arrays are shown as empty, before the state of P2 for the each time step is transferred into the associated seqCap array. After the transfer each seqCap activation becomes identical to the P2 state shown above it. That state is not shown until the next pass, when it interacts with the inputs to P2 from P1.

**Figure 2(b):**
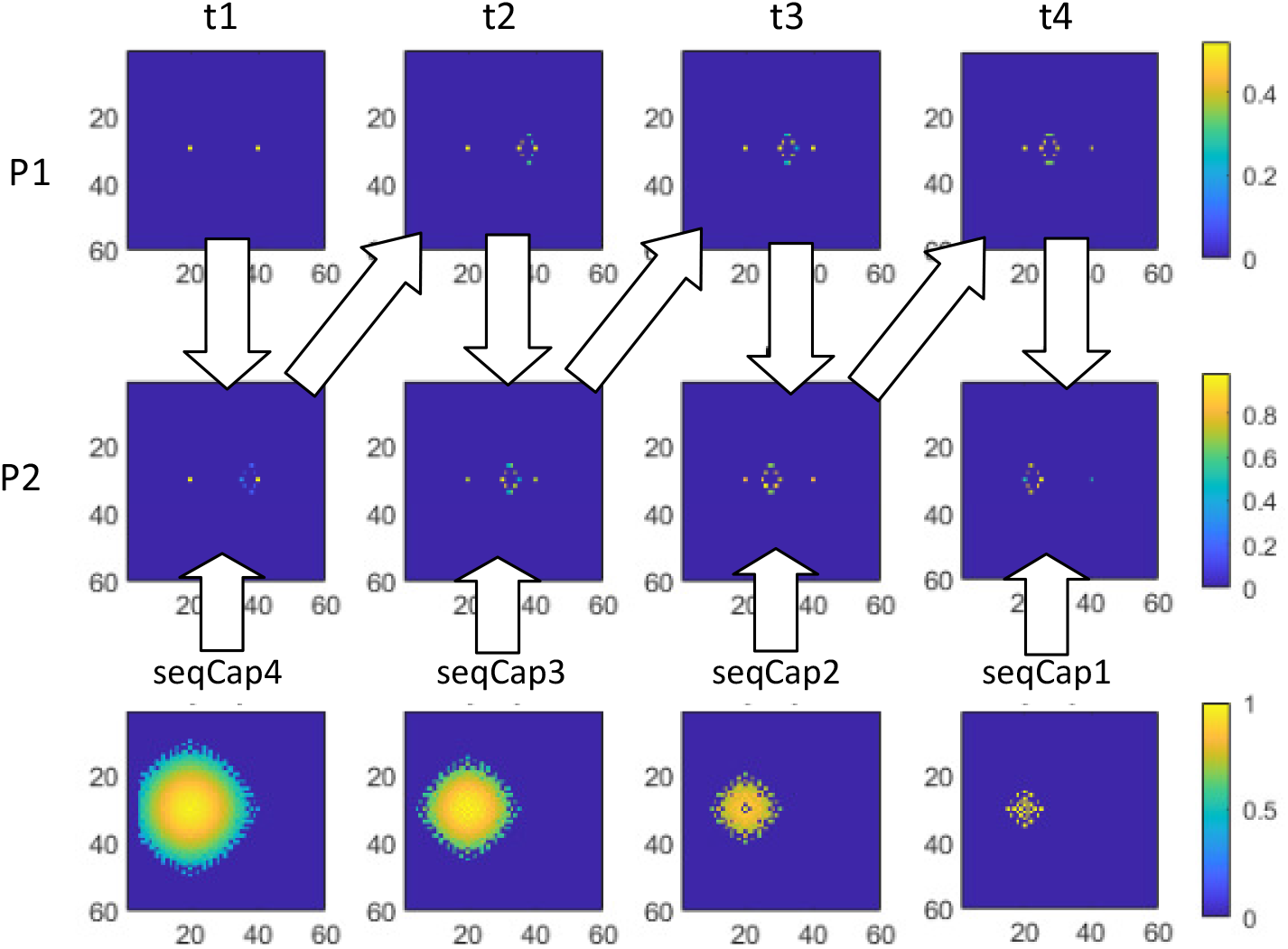
1^st^ pass backward. The activation pattern stored in the seqCap arrays which were set in the first forward pass now interact with the activations from P1 in the dendrites of P2, as indicated by the up arrows. Not shown here is the propagation of the signal from the axons of P2 at each time step to reset the stored pattern in the associated seqCap array, which will be used in the next, forward, pass. Notice the reversal of order of the utilization of the seqCap arrays relative to the previous, forward, pass.

**Figure 2(c):**
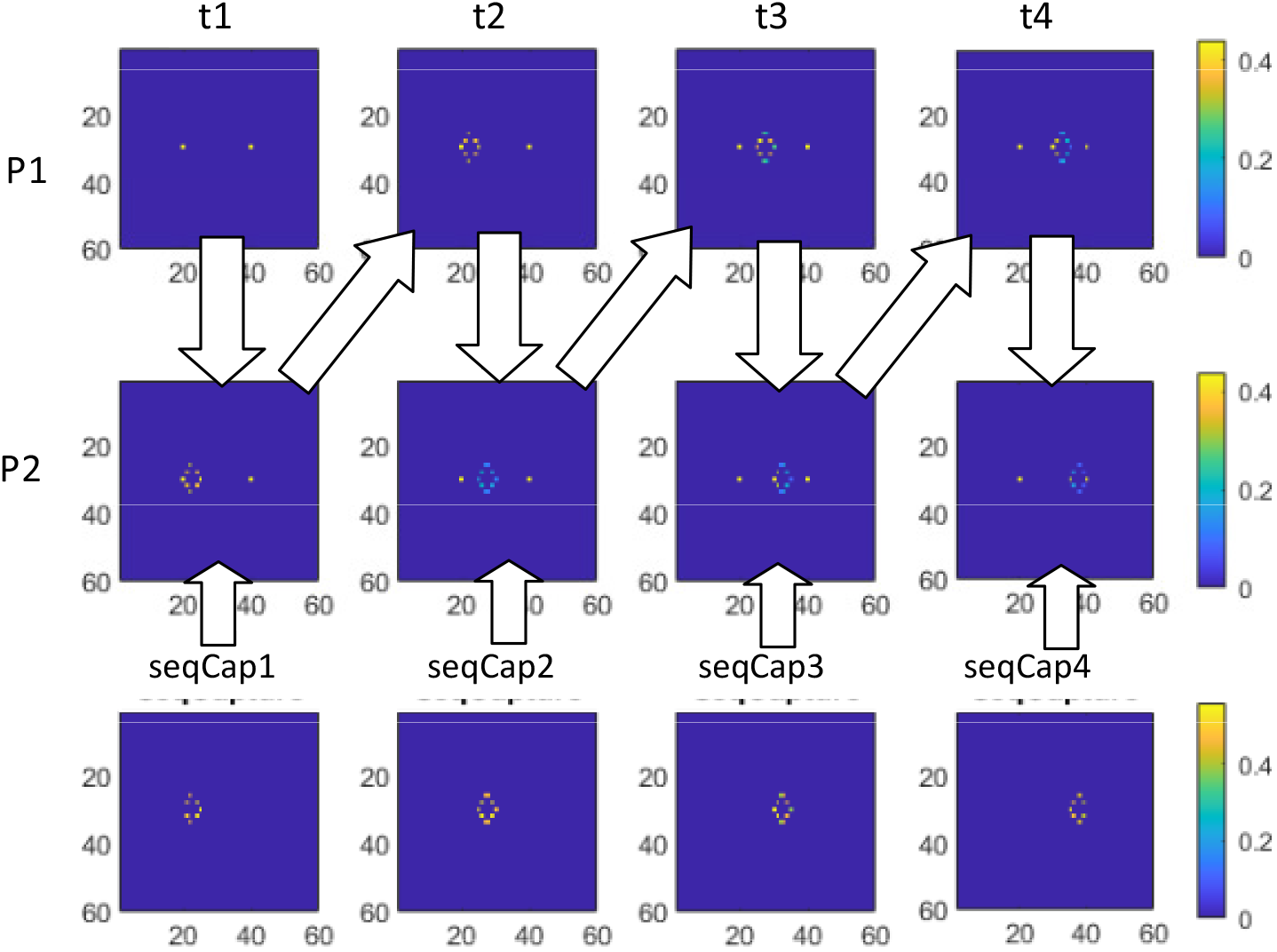
2^nd^ pass forward

**Figure 2(d):**
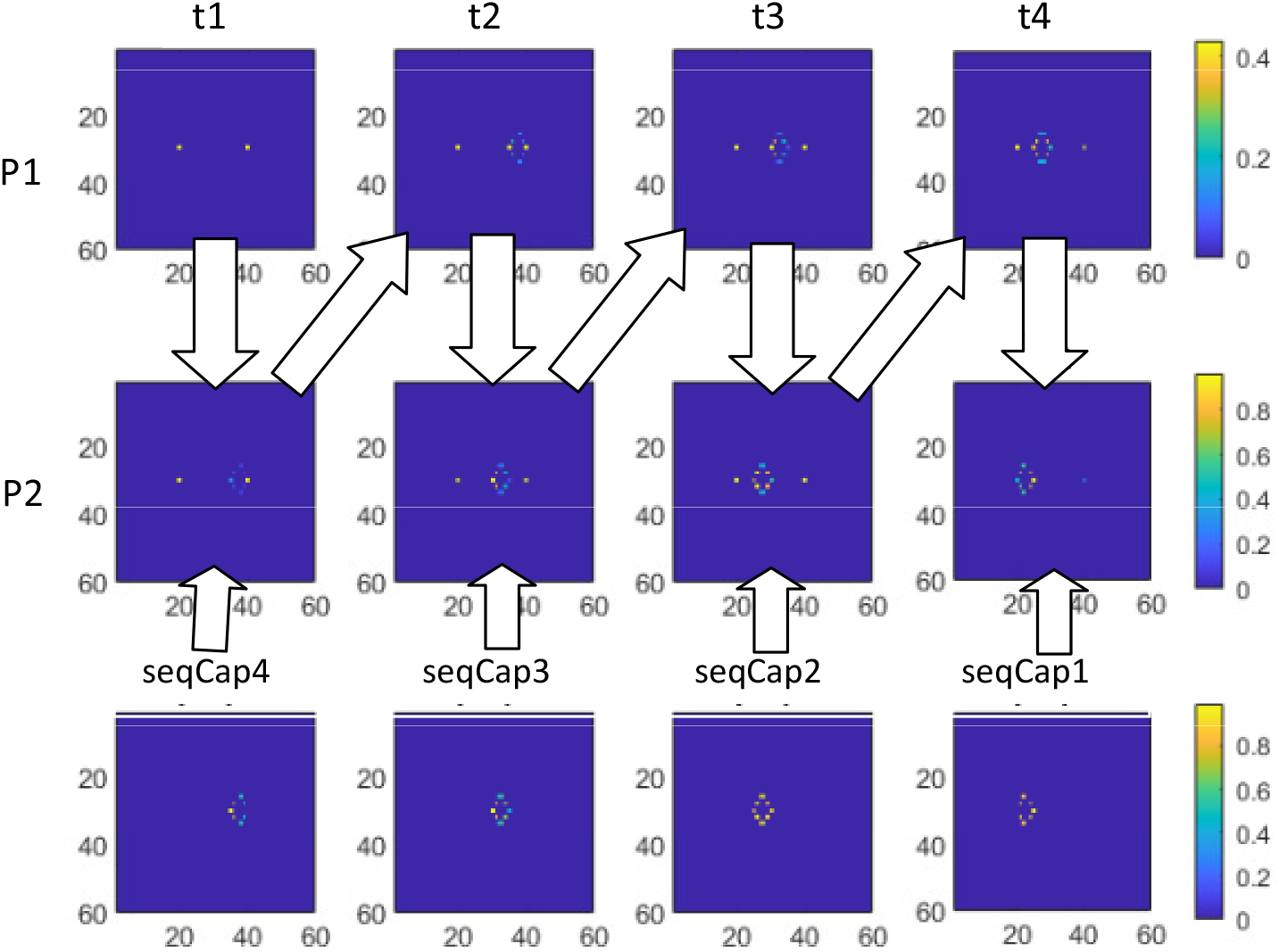
2^nd^ pass backward

**Figure 2(e):**
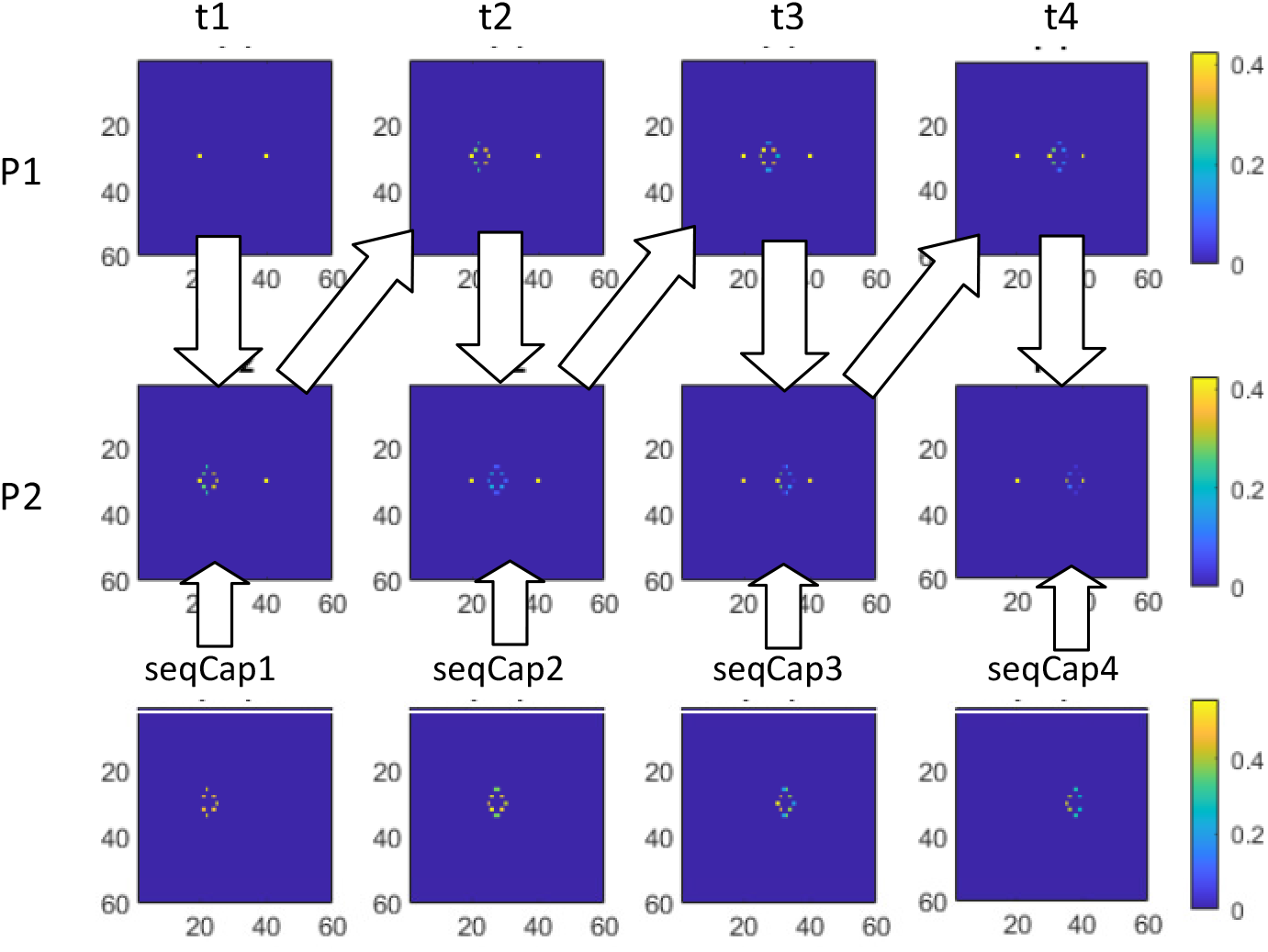
3^rd^ pass forward

**Figure 2(f):**
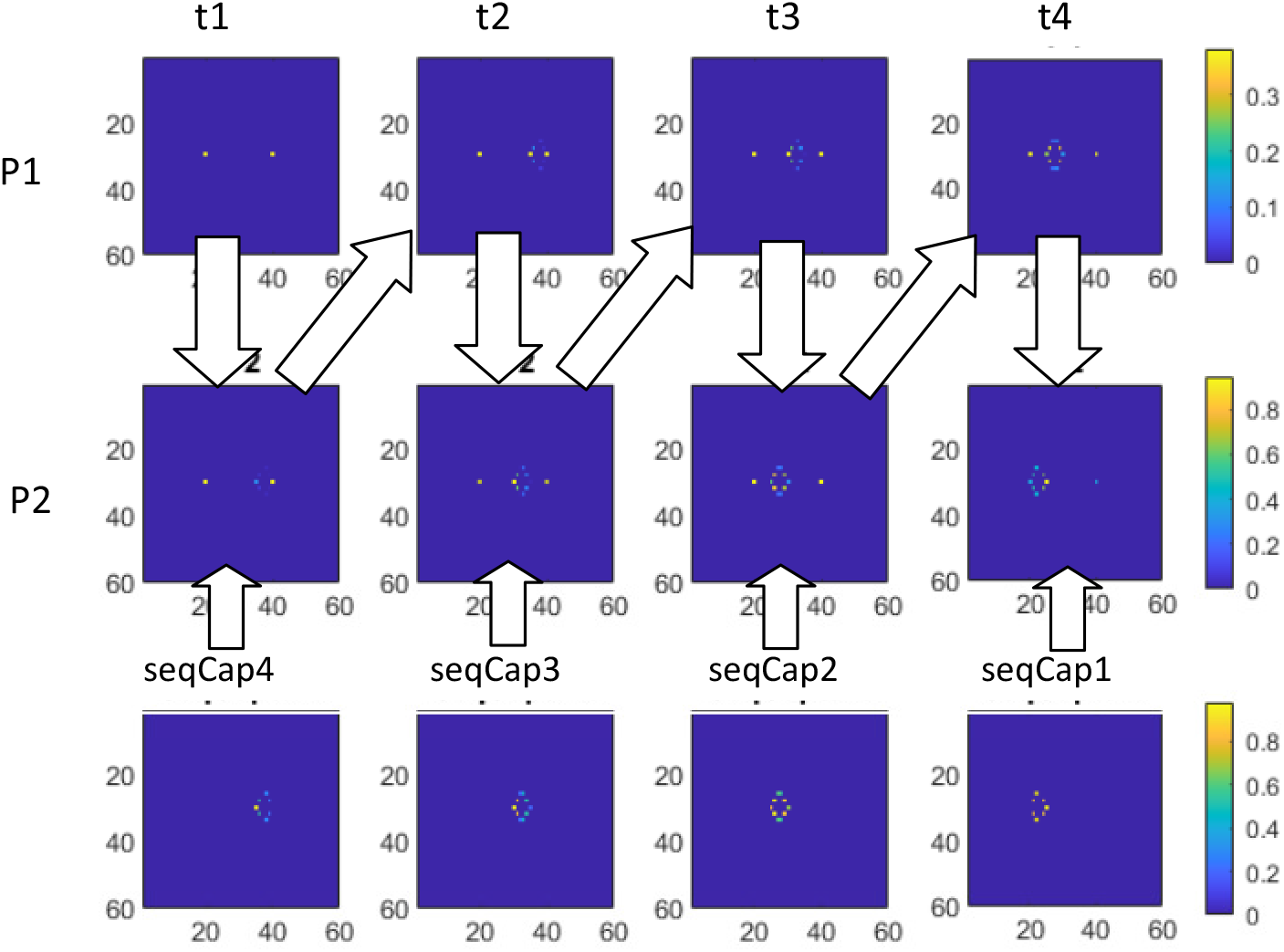
3^rd^ pass backward

The activity patterns displayed arise from the mapping connectivity pattern of the axons of P1 to the dendrites of P2. These implement a set of connections from each cell in P1 to a cluster of 28 cells in P2 radially surrounding, but not including, a central P2 with the same coordinates as the source cell in P1. The radii range from 1 to 5 place cell loci. These mappings are responsible for the steps in the path to be generated. The specific parameters of the mappings used in the simulation are presented later.

SeqCap is the component of the architecture responsible for the convergence of the computation to a solution. It will be described functionally at first, since there are at least two kinds of possible implementation which are computationally equivalent. SeqCap is a structure which can in effect take a series of snapshots of the pattern of activity on all the axons of P2 at a particular time and recall these activity snapshots on demand after some short interval of time. The recall of each snapshot reproduces the activity pattern of P2 at the time the snapshot was taken, and uses it as an input to the dendrites of P2 at a later time. The details of how seqCap stores its series of snapshots and how it reproduces the pattern of P2 activity on demand is not important in understanding how the computation takes place. But the likely neural implementation, which addresses function in more detail, will be discussed in detail later.

In the toy problem below one can conceptualize SeqCap to be structured as four separate 60×60 cell arrays, each of which can hold the entire state of P2 axons at one step of the computation and feed it back to P2 dendrites during the subsequent pass. The details of this will become apparent in the animation that follows.

In the simulation displays of Figure 2(a-f) time proceeds to the right, four time steps. Each panel of 12 graphs represents the progress of the computation in one direction, i.e. one pass. Time is always represented moving to the right.

In the simulation the value of each pixel, as color coded, represents the activation of each cell in the array. For simplicity non-activation will be referred to as 0.0 and full activation will be referred to as 1.0.

The first pass, as indicated in Figure 2(a) is always forward, that is, proceeding from the path start location. In Figure 2(a-f) the start location is at coordinates (20,30). The goal location is (40, 30). These are highlighted for reference, but do not necessarily represent activation.

In Figure 2(a) the bright spot at the start location for the P1:t1 frame is coincident with the initialization activation. The bright spot at the goal location in all P2 frames identifies the place cell that the goal-detect cell will be triggered by during the forward pass if it becomes active, but that bright spot does not indicate activity. (In frame P2:t4 the locus of the goal-detect trigger is identified by an orange arrow.)

#### The computation process for the first step of the first forward pass, t1

At the beginning of the process the entire place cell arrays P1 and P2 are made inactive (i.e. set to 0.0) by general inhibition. The place cell at the start location in P1 is then set active (i.e. set to 1.0) via the initialization inputs (probably from EC) shown in Figure 1.

The mapping connectivity from P1 to P2 spreads the activation to 28 cells in P2 at radii from 1 to 5 centered on the locus of the active cell in P1. The central cell of this pattern in P2 is not activated. The P2 activation pattern is then transferred back to P1, as indicated by the arrows. (*This direct connection which takes P2 output to P1 input is probably not neuroanatomically accurate, as most accounts put the feedback through subiculum. There is, however, evidence that the simplification is functionally correct. This will be discussed later*.)

The pattern of activation from the P2 axons also flows into seqCap1, one array of the four in seqCap, where it will be held until the next pass. As noted in the Figure 2(a) captions, this state of seqCap1 is not displayed until the following pass, in this case Figure 2(b), when it is used as part of the computation. Consequently, in Figure 2(b) below, the stored pattern in the seqCap1 array is the same as the first (leftmost) activation of P2 in Figure 2(a).

#### The computation process for the rest of the first forward pass, t2-t4

At each time step the axon of each active cell in P1 activates the dendrites of a group of cells in P2 determined by the mapping connectivity pattern. Because a pattern of radius 5 in P1 has been received from the first step, the mapping of activity to P2 now spreads the radius of activated cells even further at each step. At each step a wider activation is passed back to P1 and transferred to the arrays seqCap2, seqCap 3 and seqCap4 in successive steps. The successive selection of seqCap arrays is controlled by a daisy chain of enabling neurons which is advanced by each wave of activation from P1 to P2. (Discussed further in the navigation circuit section below and illustrated in Figure 9.)

In time step t4 the spread of the activation in P2 has now reached far enough so that one activated cell will be at the locus of the goal. This state is sensed by a goal-detection neuron (or more likely group of neurons) whose dendrite spans the axons of P2 and has a strong synapse with the P2 cell corresponding to the goal locus. (How this is initialized will be discussed in the circuit section below. See Figure 9.) Once activation reaches the goal and the goal-detection neuron fires, the first forward pass is complete. The firing of the goal-detection neuron(s) triggers the re-initialization of P1 and P2 as described shortly, and triggers the first neuron in a seqCap selection daisy-chain opposite the one sequencing the forward pass (Figure 9). The first backward pass is thereby initiated, and continues as will be described shortly.

In this toy problem the distance between start and goal locations is 20, and the maximum radius of the mappings is 5, so takes 4 steps for the activation to reach the goal. The symmetry of the increase in the activated region insures that the first contact with the target location takes place after the minimum number of steps, and ultimately results in a near optimal path.

#### Computation process for the first step of the first backward pass

Instead of labeling the next step t5, for clarity the steps of each pass will be labeled t1-t4 because it makes the utilization of the seqCap arrays clear.

On the backward pass the cell arrays P1 and P2 are set to 0.0 by inhibition and then the cell in P1 at the location of the ***goal*** is set to 1.0 via the initialization input to P1. (This input also likely originates in EC.)

As for the forward pass, the active cell in P1 is mapped to the dendrites of the 28 surrounding cells in P2. The same mapping pattern is used as in the forward pass. But starting with this pass there is an additional input to the P2 cells from the stored pattern in seqCap4 (which has been set in the previous, forward pass). On backward passes seqCap arrays are selected in the opposite order from that of the forward passes. The successive selection of seqCap arrays is controlled by the aforementioned daisy chain of enabling neurons coupled in the opposite order from those engaged during the forward pass (Figure 9). Again, the daisy chain activation is advanced by each wave of activation from P1 to P2.

The input from seqCap to the P2 dendrites comes through some intervening microcircuitry (one possible version discussed in the section on circuits) to the P2 dendrites, proximal to the synapses of the mappings from P1. The microcircuitry “inverts” the signals so that the 0.0 outputs inhibit (and the 1.0 outputs do not inhibit) the distal signals in the associated dendrites of P2. The net effect is that the stored pattern in seqCap4 “gates” the distal signals such that **only those cells of P2 which receive inputs from P1 mappings AND seqCap4 become active**.

This gating is referred to as ***intersection***, as in the set operation which selects the elements common to two sets. In other words, the pattern of activation from P1 mappings is filtered by the pattern of activation in seqCap4. This filtered pattern becomes the pattern of P2 axon activation. This pattern is now transferred to the dendrites of P1, and to the seqCap4 array, where it replaces the stored pattern acquired in the first forward pass. This new pattern is only displayed in the seqCap panel for the subsequent, forward, pass, when it becomes part of the intersection operation.

The effect of the intersection operation becomes evident in Figure 2(b) time step t1. Without the intersection operation the activation pattern in P2 would densely surround the goal locus to a radius of 5. It would look much as P2 in time step t1 in Figure 2(a) except at the goal end. Instead, the activation pattern in P2 has only a few active cells, and they are clustered about 5 pixels to the left of the goal locus. This is because only non-zero loci at the very periphery of the large activation in seqCap4 are co-located with the P2 cells which have received activation from the radius of mappings from the single active goal cell of P1 in t1, and the intersection operation results in the selective reduction.

#### Computation Process the rest of the first backward pass, t2-t4

Again, the axon of each active cell in P1 activates the dendrites of a group of cells in P2 determined by the mapping connectivity pattern. The tight cluster in P1 received from the first step is mapped to a broader radius in P2. But this wider activation is now intersected with, or gated by, the stored pattern in seqCap3. The result is the small cluster of active P2 cells seen at t2. This cluster of activation is now transferred seqCap3, replacing the earlier pattern. It is also passed back to P1 for processing in time step t3.

The same sequence takes place in t3 and t4. It is now apparent that the progression of seqCap cell arrays (seqCap4, seqCap3,…) used to gate the mappings from P1 goes in the opposite order to the time sequence t1, t2,… This is true of all backward passes. Also, the activity patterns in P2 replace the patterns in seqCap2 and seqCap1 in t3 and t4, respectively.

The furthest extent of activation from either the start or goal loci can be thought of as the activation wavefront. The path reduction process is therefore the result of the intersection of the forward and the backward wavefronts. At each step the radius of the forward activation from the start location plus the radius of the backward activation from the goal location is greater by up to 4 than the distance between the start and goal loci. Where they overlap is the surviving path.

Unlike the forward pass, what stops the backward pass sequence is that seqCap1 is used for the last gating and then for storing the state of P2 axons in time period t4. This stops the backward pass, and the second forward pass is initiated. As seen in Figure 2(b) this has brought the steps of the path back to the start location.

#### Computation Process for Second Forward Pass, t1-t4

Again the P1 and P2 arrays are initialized as they were in the first forward pass. The second forward pass is computed as the first forward pass, except that in this and subsequent forward passes the mappings of activity from P1 to P2 are gated by the stored patterns in seqCap1 through seqCap4 in order. So, in forward passes the seqCap arrays are utilized for gating in the forward order, in contrast to their utilization on backward passes. The activity of the P1 and P2 arrays are seen in Figure 2(c). And the activation pattern of P2 is again stored in one of seqCap1 – seqCap4 in order, but not displayed until the next pass, Figure 2(d).

The second and subsequent forward passes are terminated by the activation in P2 reaching the goal cell location, as occurred in the first forward pass.

#### Process for Second Backward Pass, t1-t4

The processing of the second backward pass is identical to that of the first backward pass.

#### Subsequent Passes

It is apparent in Figures 2(c-f) that the patterns of activation don’t change much. In practice full convergence is reached by the end of the second forward pass. The activations of P2, as captured in seqCap can be narrowed by selecting the most active cells at each step. The resulting paths are displayed in Figure 2(g-i) for three iterations, that is, three pairs of forward and backward passes.

**Figure 2(g-i):**
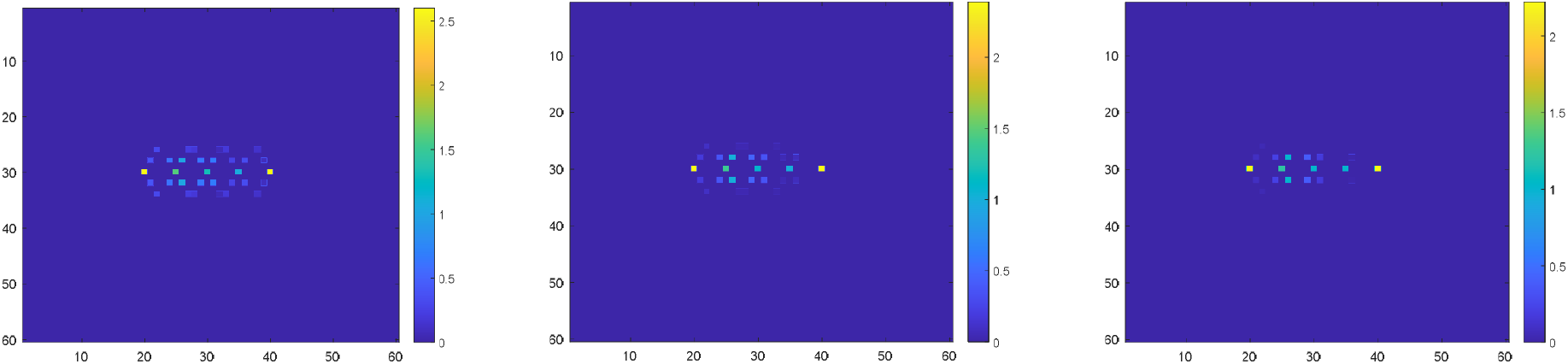
Sequence capture linear after 1^st^, 2^nd^, 3^rd^ backward passes, start and goal superimposed

#### Playback

Note that the mechanics of iterations 2 and 3 are exactly those which would produce the observed forward and backward playback sequences of already established (i.e. captured earlier) paths. [3] In this case both the P1 start or goal locus initialization and the contents of seqCap would be provided by longer term memory holding the previously experience path.

Figure 2(g-i) shows a representation of all of the SeqCap arrays combined into one, forming a time series display of the surviving path activations at each iteration. The start and goal loci are displayed bright. It can be seen that only the direct path between the start and goal place cells have maximum activation, while those to the side of that path have activations rapidly decreasing. Thresholding or local winnertake-all would reduce the path to a sequence of single active cells. Figure 2(g-i) makes it clear that the convergence is effectively complete after the first backward pass.

#### Mapping

In all the navigation simulations in this paper the mapping operation is implemented as a set of offsets to compute the destination locus in P2 from the source locus in P1. In each step each active cell in P1 is activates a the dendrite of a cell in P2 at a direction and distance specified by one of 28 mapping vectors in the following list:

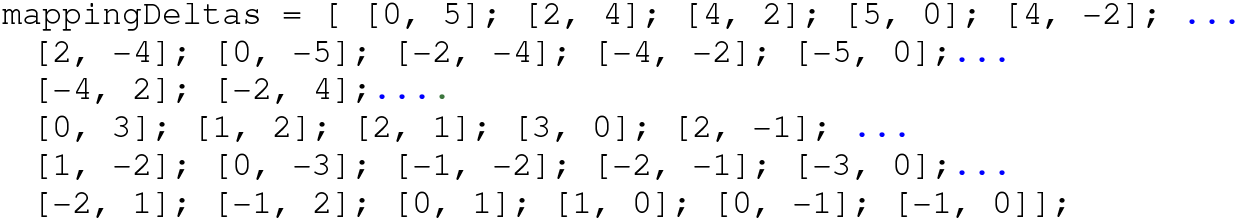

In other words, if the coordinates of an active cell in P1 are (x,y), the cells in P2 are activated at locations (x+dx, y+dy) for each [dx, dy] pair in the mappingDelta list above. If the same destination cell is activated by more than one mapping, all activation contributions to that cell are added, thus forming a *superposition* of activations.

Since the maximum mapping distance is 5, for the short distance between start and goal in Figure Z(a-f), it takes four steps to reach from start to goal place cells

#### Superpositions

In Figure 2(a) it is apparent from the color coding of activations that the activations after the first step are not constant across the diameter of the active patch. The activations toward the center are stronger. This is because each active cell in P1 is mapped to 28 surrounding cells in P2, so the cells toward the center receive more input than those on the periphery. Since the activations are additive, this creates a superposition of activations.

Note, for computational simplicity, the outputs of P1 and P2 are scaled on each step, so that no cell has an output greater than 1.0. In order to make the full radius of the activation visible in the Figures, the range of activations is greatly compressed when displayed.

#### More Realistic Path Planning Problems

Figures 3(a-l) illustrate the same direct path computation as in the toy problem above, but for a more realistic length journey. The cell arrays have been enlarged to 80×80 and the start and goal loci have been spread further apart and aligned at an angle to the grid axes. Otherwise the execution remains unchanged, though of course more steps are required before the goal is reached. The operation at this more realistic scale exhibits the same dynamics as the “toy” problem above. Figure 4(a-c) display all steps after first, second and third backward passes. These are thresholded P2 activations, selecting the activations greater than 0.9 times the maximum in each step. There is some progress in convergence after the first backward pass.

**Figure 3(a-d):**
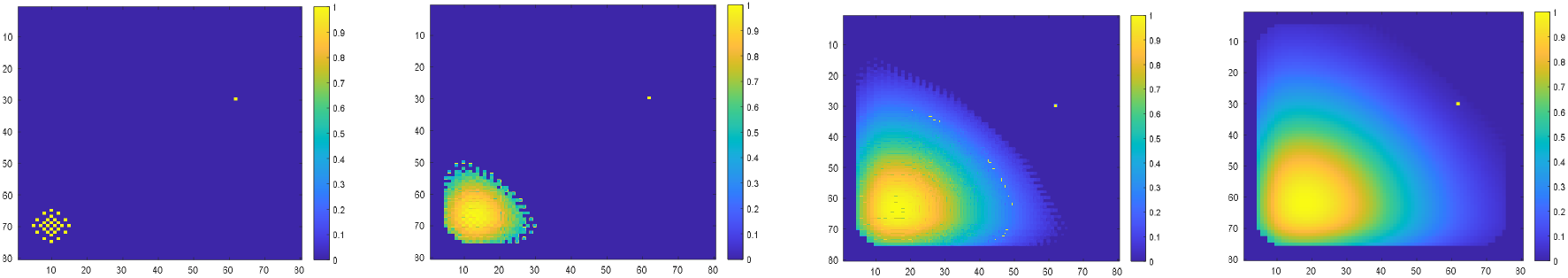
P2 activation during first forward pass. Approximately every 5^th^ step shown.

**Figure 3(e-h):**
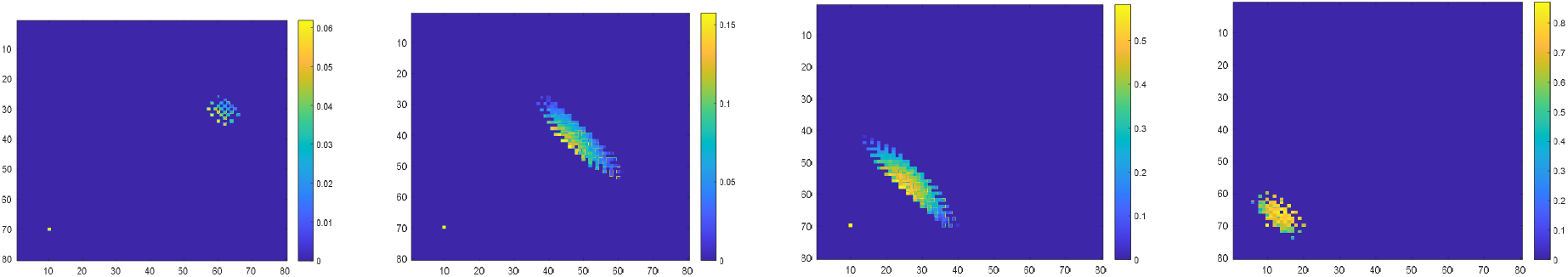
P2 activation during first backward pass. Approximately every 5^th^ step shown.

**Figure 3(i-l):**
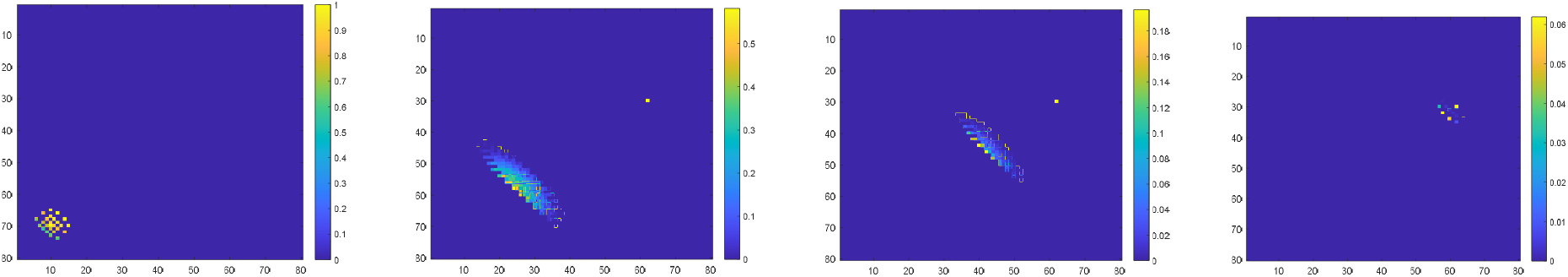
P2 activation during second forward pass. Approximately every 5^th^ step shown.

**Figure 4(a-c):**
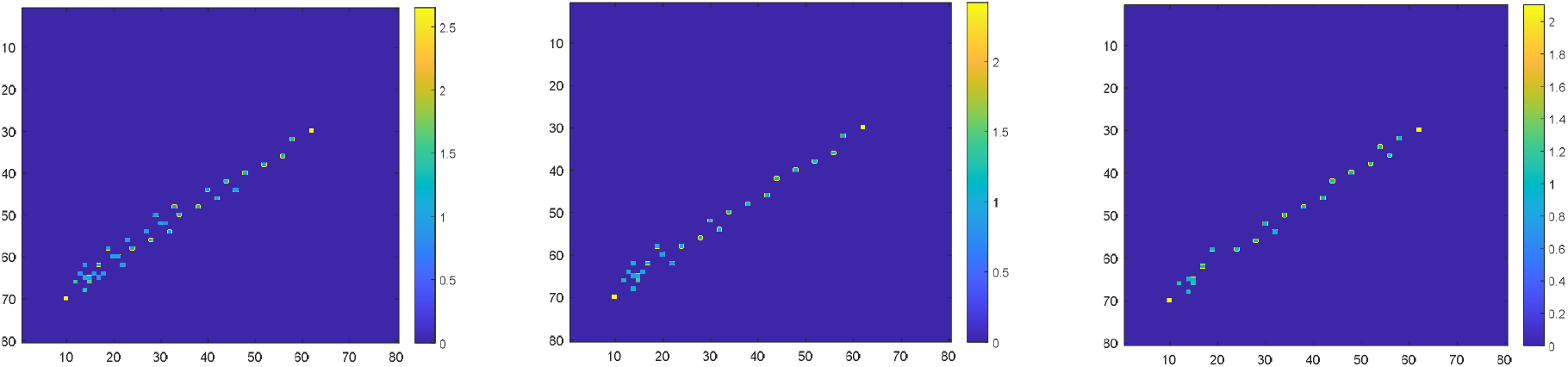
Thresholded P2 sequence for first, second and third iteration. Every step shown.

#### Obstacles in the Place Cell Space

Obstacles to the path are introduced to the place cell space by a “masking” operation which inhibits dendritic input to P2 place cell somas, thus excluding certain P2 cells from being utilized in the path solution. In the simulation an array of the same dimensions as P1 and P2 contains non-zero values in locations where the place cell space is to be excluded. In biological terms, the masking operation would be the mechanism by which boundary cells [7] preclude the Hc path solution from crossing the “protected” boundaries. In the simulations below the entire excluded area is masked, but masking just the boundary around the exclusion zone suffices. So long as the masked boundary region is wider than the longest mapping radius, the path cannot cross into the excluded area. When an obstacle is represented by an exclusion zone in the place cell space, the solution “cloud” extends around that obstacle until it reaches the goal locus on the first forward pass, as seen in Figure 5(a,b). Figure 5(c-f) displays four snapshots of P2 activity on the backward pass, skirting the obstruction. The intersection with the forward pass activations has narrowed the path significantly.

**Figure 5(a,b):**
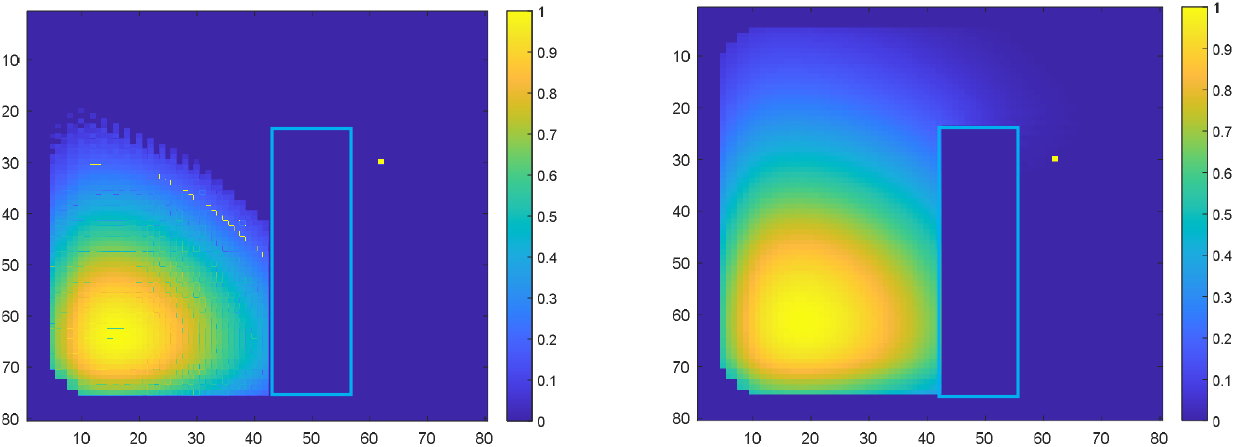
P2 first forward pass, two snapshots. Obstacle boundary highlighted.

**Figure 5(c-f):**
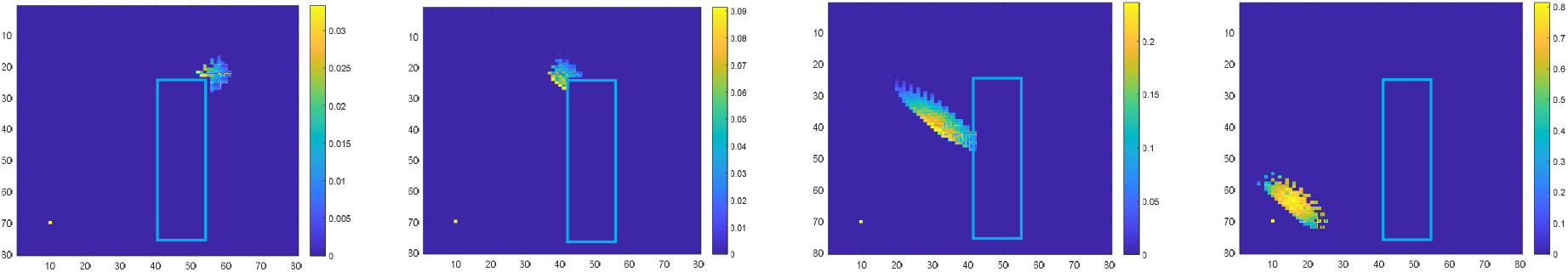
P2 first backward pass, four snapshots. Obstacle boundary highlighted.

The completed path is rendered in Figure 6(a-c) by superimposing all SeqCap arrays after each backward pass. The region of the obstacle is made visible by subtracting *obstMask,* i.e subtracting 1.0 in the locations of the obstacle. The result is thresholded to reduce the path to the most strongly activated place cells at each step. By the end of the second iteration the convergence has reached its fixed point.

**Figure 6(a-c):**
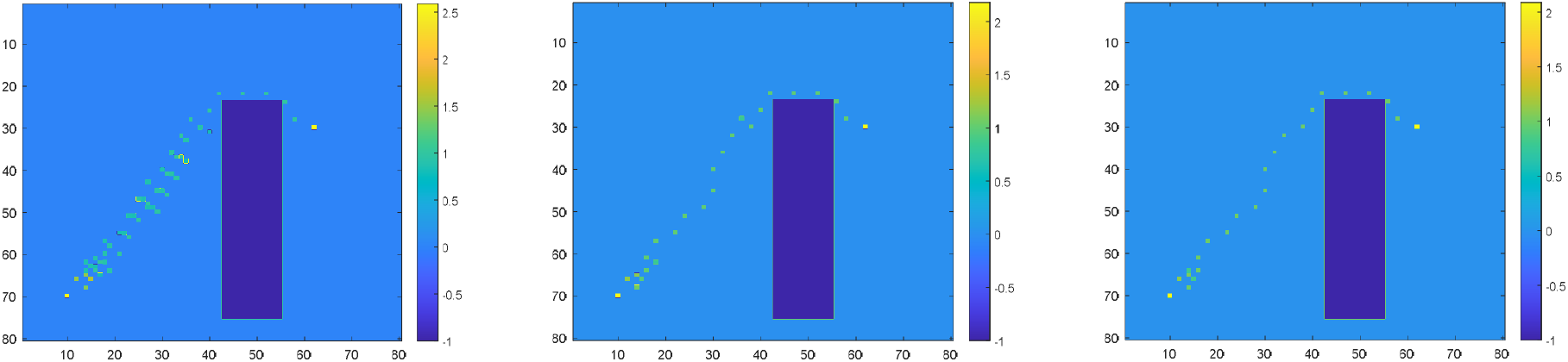
P2 sequence thresholded sequence for first, second and third iteration

Figure 7(a,b) shows allowed areas corresponding to streets in a city neighborhood (in London) along which the SSC finds the shortest path between start (lower left) and goal (upper right) constrained to follow the city streets. In this case the masking elements are set to 1.0 for all excluded areas, and to 0.0 for the permitted street areas. This demonstration is motivated by the association between the famous path solving capabilities of London taxi drivers and the observed unusual Hc development of those drivers. [8]

**Figure 7(a,b):**
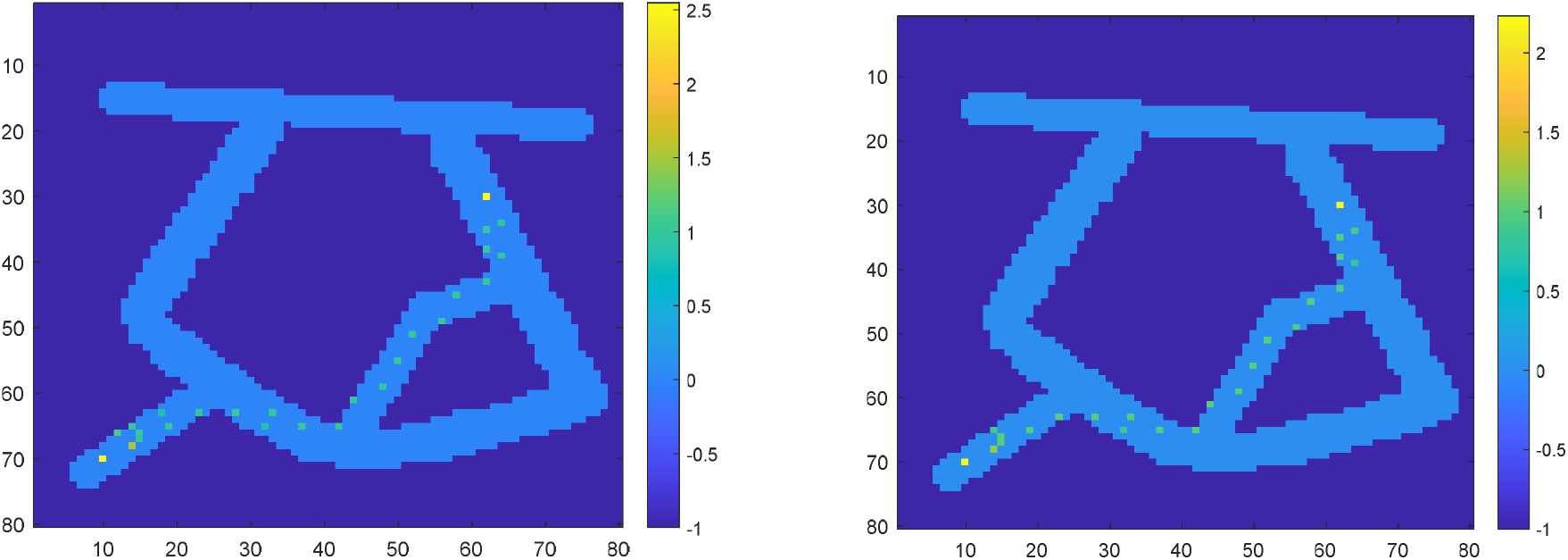
P2 sequence thresholded for first and second iteration: city mask

#### Characteristics of the Process

The exposition of the algorithm in “program” form, Appendix 1, makes it clear that weight updating plays no part in the pathfinding operation. This avoids any biologically questionable reliance on deltas of high precision to implement an update. In this respect the algorithm has no relationship with neural nets. (This characteristic may also have been noted by the reader in the earlier narration of the process.) The dynamics of process presented above emerge from signal reinforcement and suppression, which is why it arrives at nearly full convergence after one iteration (i.e. one forward and one backward pass).

Reinforcement and suppression are operations requiring minimal precision and are entirely plausible for the known repertoire of neuronal signal interactions.

Hc is often ascribed the capability to compute “path integration” or, to use the more precise robotic term, odometry. This is the use of knowledge of heading direction and speed of travel to compute current location given knowledge of the starting location. It is a “forward” computation related to the “inverse” computation of path finding. It requires the mapping vector to be selected by knowledge of heading and speed. The neuronal circuit equivalent of the path finding algorithm (discussed in a later section) requires only a minor addition of selectable mappings and thereby become capable of path integration. This is discussed in Appendix 2.

#### Relationship Computations

The second major set of Hc functions, as mentioned in the Introduction, are those associated with the implementation of cognitive maps. A very simple extension of the algorithm (and circuitry) described above can enable it to establish pathways through cognitive maps as well as through the physical place maps as demonstrated in the preceding sections. Assume that two place cells, A and B, are randomly assigned by the lateral entorhinal cortex (LEC) to represent two concepts or items. An ordered relationship between the actual entities represented by A and B, such as the transitive relation A > B, can be established by a one way journey from place cell A to place cell B. This geometry is in effect a directed graph which can represent preference order, temporal order, causal order, etc. The mechanism already described for navigation can establish and memorize a sequence of short steps in a more or less straight line connecting locus A to locus B.

Assume another place cell is assigned to concept or item C, and the relationship of the entities C and B represent is known to be B > C. Since the locus for B is already established, the navigation mechanism can readily find and memorize the directional path from locus B to locus C.

Now if one wants to discover if there is an inferable relationship between A and C, the existence or non-existence of a sequence of memorized paths between A and C can answer the question. In the situation described, there are memorized paths from A to B, and from B to C, so the relationship A > C can be validated. The navigation algorithm described earlier needs a small bit of extra machinery to determine if a useful sequence of memorized paths exists. As described above it can’t do that on its own.

But before describing that extra machinery, a problem with using “pure” path navigation, as described above, needs to be pointed out. In traversing the sequence of place cells from A to B, for example, a place cell assigned by LEC to some other entity may be inadvertently activated. This brings another entity into the relationship by accident. That may or may not have disruptive consequences. It would be far safer (and faster) to make the journey in one leap from A to B without treading on any place cells in between.

The single-leap journey can be effected by a modification to the mapping step in the algorithm presented in the earlier navigation discussion. Instead of the mappings being fixed short offsets or deltas from a location of origin, the mappings will be memorized pairs: an origin locus and a destination locus. So, in the above example, when the relationship A > B is declared, a pair of locus-specifying cells or arrays become associated: the first activating the place cell representing A, and the second activating the place cell representing B. It will be shown later that this can be implemented by a single or a few neurons, but for conceptual simplicity it will be assumed that A and B are each specified by an array of cells conforming one-to-one with P1 and P2. In these arrays only one cell is non-zero, the one corresponding to the place cell loci of A or B.

Assume that the array pairs {A B} and {B C} have been established as just described. Now to test for the existence of a relationship journey from A to C, it is only necessary initialize the A location in P1 and set the goal-detecting test on the output of P2 to trigger when the C locus is active. The mapping mechanism will then grab all the array pairs whose first member is A, and use the second element of each pair as destinations of the mapping into P2. One of these destinations will be the place cell locus for B and there may be others. That total activation of P2 will be transferred into P1. The mapping mechanism will now activate in P2 the locus of the second entity of any pair whose first element is activated in P1. Since the place cell for B is active in P1, the array pair {B C} will be invoked and the place cell for C will be activated in P2. This will trigger the goal-detected test, establishing in two cycles that a path from A to C exists.

Now there may be many other place cells active as well, so in order to prune the set of active place cells to just the path A > B > C the process described needs to be run backward, just as in the navigation case.

This process is better illustrated by a more complex case, which contains second order transitive relationships and several distractor entities.

For the demonstration arbitrary spatial assignments of place cells represent entities A-H, as in Figure 8(a). In this use of the place cell space there is no meaning to any spatial order. A different graphical representation is used here as a result of the random spatial assignment to entities. Having separate panels for P1 and P2 makes the association between related pairs of entities very difficult to visualize, as the two loci for each relationship appear in different panels. Instead, the relationships are made explicit by displaying both loci for all pairs in a single panel with lines connecting the two loci of a given pair.

**Figure 8(a):**
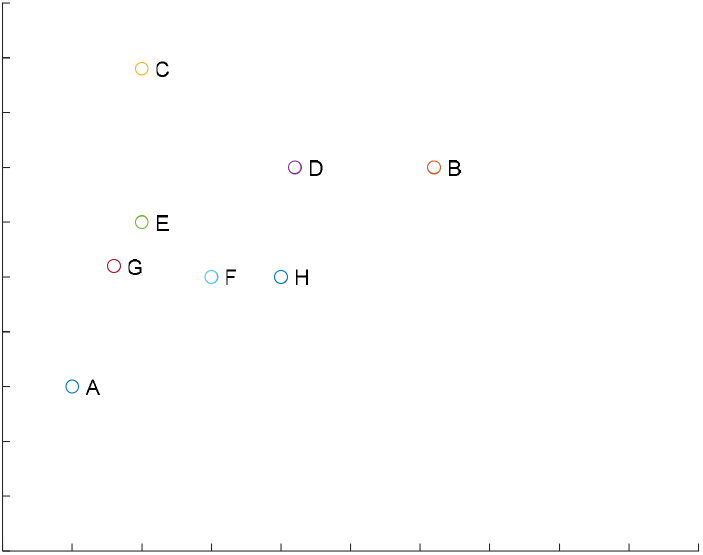
random assignment of place cell loci to non-spatial entities

##### Setup

The relationship pairs are established. These define the available set of mappings between the locus of first entity of the pair and the locus of the second entity of the pair: e.g. locA is the coordinate pair (Ax, Ay) in the place cell space, etc.

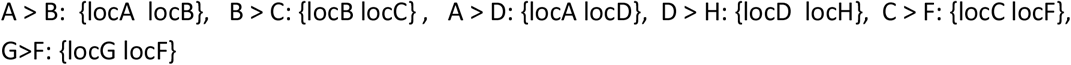

Establish the relationship under test: A > F? The place cell locus for entity A is defined to be the start of the “journey” and the locus for F is defined to be the goal of the journey.

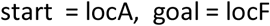

##### First forward mapping step

The start place cell at locA is initialized to 1.0 in P1. All established mapping relationships in the list above whose first element is locA become active and activate in P2 the cell specified by the paired second locus. Thus the cells in P2 which become active represent all entities with direct relationships with A. The Figure 8(b) shows the mapping of loci representing entities. In this case A is in P1 and B, D are in P2. B and D are also captured in seqCap1.

**Figure 8(b):**
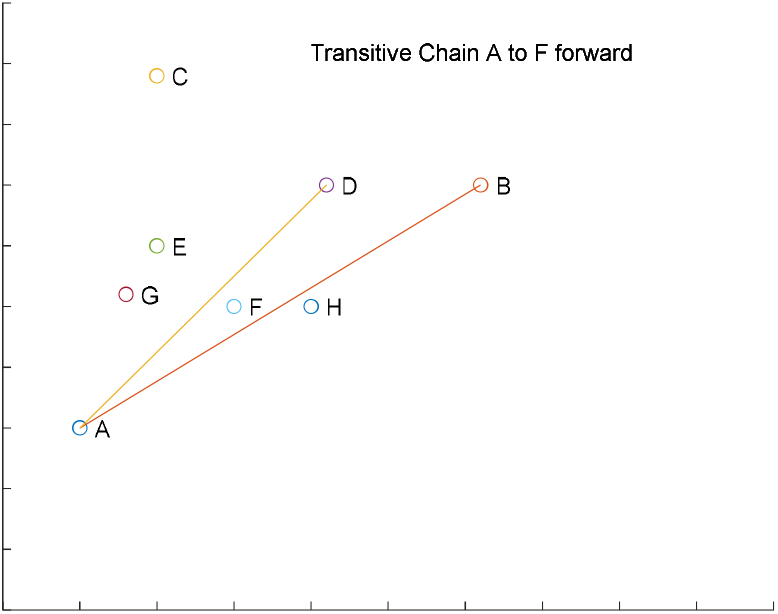
first forward mapping, from locus of A

**Figure 8(c):**
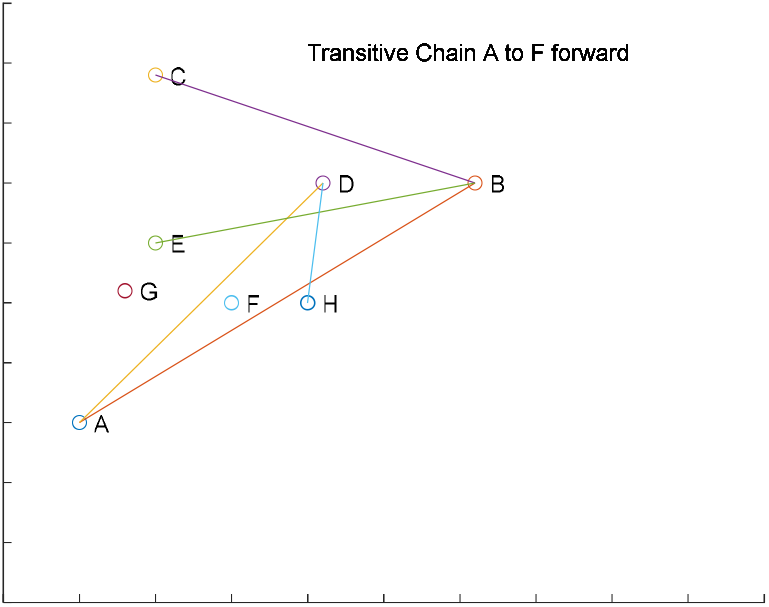
second forward mapping, from loci B and D

**Figure 8(d):**
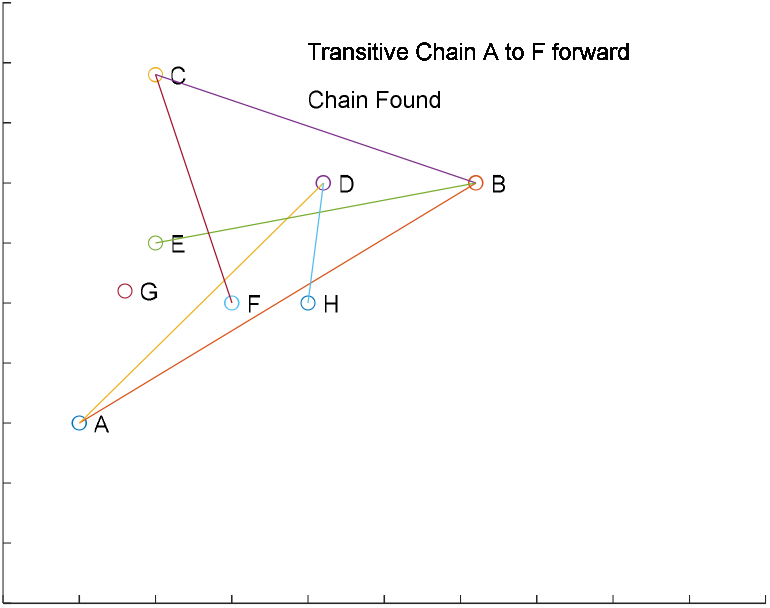
third forward mapping, from locus of C

**Figure 8(e):**
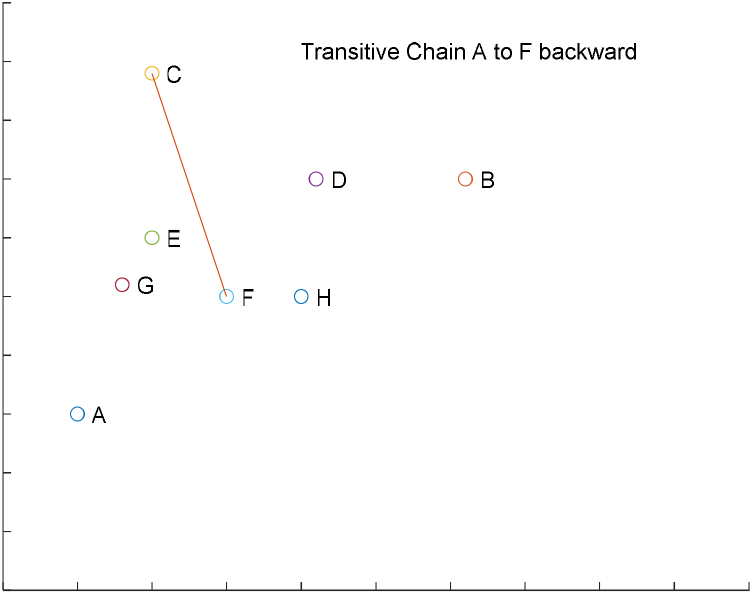
first backward mapping, from locus of F

**Figure 8(f):**
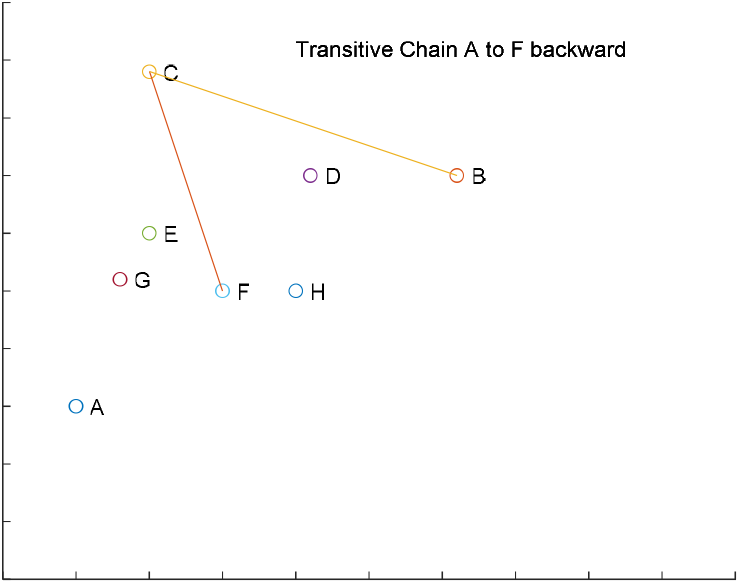
second backward mapping, from locus of C

##### Second forward mapping step

When P2 activation is moved to P1, B and D become active in P1. Now all mapping relationships whose first elements are either locB or locD are activated and their associated targets, locC, locE and locH, become active in P2. C, E and H are captured in seqCap2.

##### Third forward mapping step

When P2 activation is moved to P1, C, E and H become active in P1. The only mapping relationships with a first element either locC, locE or locH is C>F. So only F is activated in P2. But locF is also the goal of the relationship under test: A > F? So the forward pass is complete and the transitive relationship A > F is “proved” by the forward pass of three steps. But the path between A and F is not isolated from all the other entity loci which have been activated along the way. (Note relation G > F has been ignored because no prior step reached locG). For problems in which isolating the intermediate steps is essential, such as determining causal chains, the state at the end of the forward pass is not acceptable because entities outside, and irrelevant to, the causal chain have been activated and not enough information is yet available to discard them.

##### First backward mapping step

The backward pass prunes ordering relationships to “critical path”, eliminating all the spurious activations. In the first backward step, locF is set as the start of the search in P1 and (unlike in the navigation application) locA is set as the goal-reached for P2. From the set of relationships above only those with its **second element** at locF are activated. There is only one: C > F. The mapping is activated backward relative to the forward pass invocation. So the **first element** from that relationship, locC, is the mapping target in P2. Because C was an activated locus in the forward pass, it is non-zero in seqCap2 (along with E and H, which are now irrelevant because they are not mapping targets in P2) and hence C remains active after the intersection of the mapping targets from P1 and the pattern of seqCap2 in the dendrite of P2.

##### Second backward mapping step

The activation of P2 is moved to P1. Only locC is active in P1. From the set of relationships only those mappings with it second element being locC are activated. There is only one: B > C. So the first element from that relationship, locB, becomes the mapping target in P2. Because locB was an activated locus in the forward pass, it is activated in seqCap1 and hence B remains active after the intersection in the dendrite of P2.

Now, suppose there had been another relationship defined in (a) with its second element being locC, say K > C. The locus for K in P2 would also be a target for the mapping, but K would not have ever been reached in the forward pass so seqCap1 would have no “memory” of it. Hence the activation of K in P2 at this step would be blocked by the intersection operation.

##### Third backward mapping step

The activation of P2 is moved to P1. Only B is active in P1. From the set of relationships only those second element is locB are activated. There is only one: A > B. So the first element from that relationship, locA, becomes the mapping target in P2. Unlike the navigation backward pass which is terminated one step earlier by the enabling of seqCap1 for intersection, this proceeds one more step for which there is no corresponding seqCap array, and hence no intersection operation. Because A was set at as the goal-reached target of the backward pass, the backward pass is complete. The pruning by the intersection operation has isolated the actual path of transitive relationships, A > B > C > F, as displayed in Figure 8(g). The mechanism is the same as the mechanism that isolated the paths in the navigation role, just using different mappings.

**Figure 8(g):**
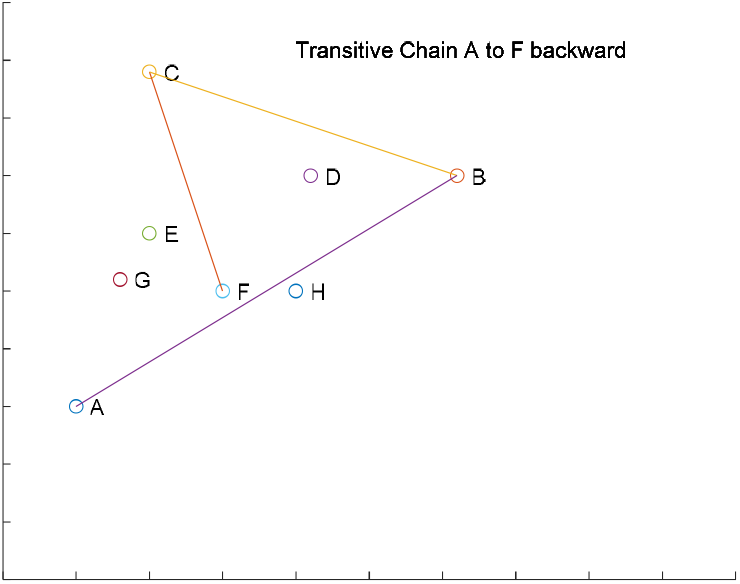
third backward mapping, from locus of B

**Figure 9:**
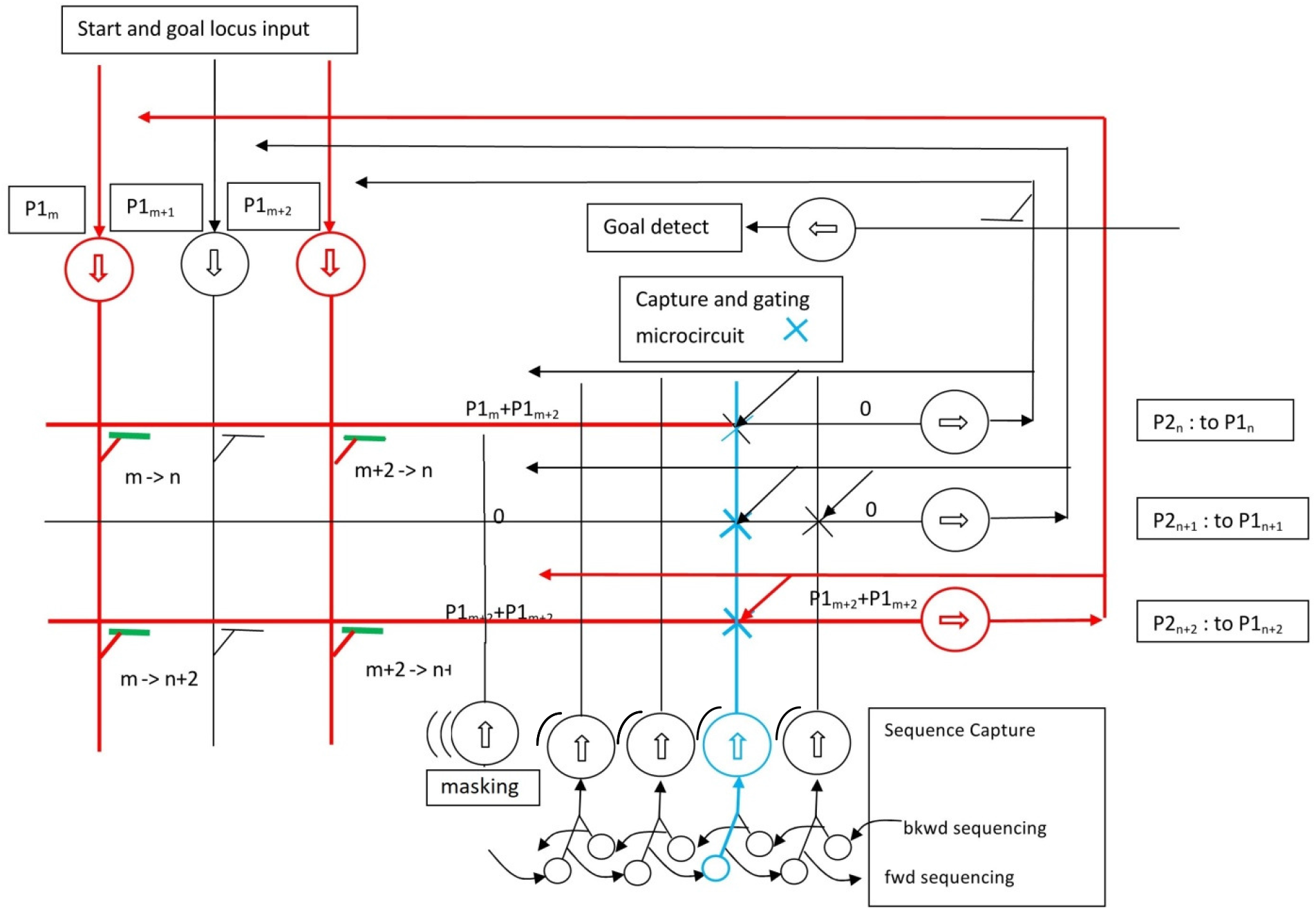
(approximate) neuronal circuit for navigational inverse problem computation. Blue X’s represent circuit details with optional implementations, see Figures 10(a,b) and 11.

The algorithmic version of the relationship computation is a minor variant of the navigation algorithm. Both appear in Appendix 1. The difference is in the mapping step.

### Neuronal Circuit Characteristics

Given the algorithmic requirements established by the simulation and some neuroanatomical and behavioral constraints from the experimental literature, it is possible to conceive an approximate neuronal circuit architecture that meets the case. For the navigational mode, such a conjecture is seen in Figure 9.

In Figure 9 the mappings from P1 to P2 are implemented by the synapses onto the P2 dendrites. Active synapses are show in green. Of course, the specific pattern of connectivity necessary to implement the direction and distance of the mappings cannot be shown with just three cells visible in each of P1 and P2. The mapping synapses between P1 and P2 are static, which is the most efficient implementation for path planning. The mappings actually used in a path solution are selected by the forward-backward signal interaction. No dynamic selection of the mappings is needed. For the other inverse problem class mentioned in the Introduction, involving higher degree relational computations, a different mapping architecture is required to implement the process illustrated in Figure 8(a-f) above. The same selection-by-intersection mechanism is used but the mappings from P1 to P2 are established differently. This additional circuitry is presented later in this section.

Because Hc is known to be involved in the inverse problem of path planning there is some debate as to whether it is also involved in the related forward problem of path integration. (Path integration refers to the computation of current location during a journey in which direction of travel is estimated from visual or other sensory cues, and speed of travel is estimated from motor signals.) This paper takes no position on that debate but in Appendix 2 discusses the addition of selectable mappings to the circuit of Figure 9 which would make it capable of path integration. The direction and span discretization of these mappings would be specific to path integration and independent of the mappings used for path planning.

In Figure 9 each seqCap array is rendered simplified as a single cell, with a graphic hint that more cells are represented. Were it possible, the rendering would show as many cells in each seqCap array as in P2, with a one-to-one connection from the axons of each seqCap array to the dendrites of P2. There are alternative architectures with much lower cell population requirements. One is discussed below.

The function of the daisy-chain of sequencing neurons has been introduced earlier. It will be recalled that a different seqCap array captures the P2 state for each step of the computation, both forward and backward. For this to take place, a daisy-chain of control neurons propagates a selection signal to enable the specific seqCap array for both the capture of the P2 axon activation pattern and the provision of the intersection pattern to the P2 dendrites. There are two daisy chains, one for forward pass sequencing and the other for backward path sequencing, as discussed earlier. The first neuron in each daisy chain is activated at the start of its respective pass, and the signal steps down the chain, advanced by the flow of activation from P1 to P2. The details of this triggering are not shown to keep the diagram relatively simple.

In Figure 9 the axon of the seqCap “cell” symbol as depicted represents as many axons as there are cells in the P2 cell array. So the X’s represent the connection circuitry from many axons, each to one P2 dendrite. There is also an input from each P2 dendrite to this circuitry designated by X. The latter enables the capture of P2 outputs to seqCap axons. Though there are several possible implementations of the part of the circuit represented by X, a detailed expansion of a likely version of seqCap and its connections to P2, requiring far fewer cells for seqCap memory than the full cell array, is presented in Figure 10(a,b) and discussed in the accompanying text.

**Figure 10(a):**
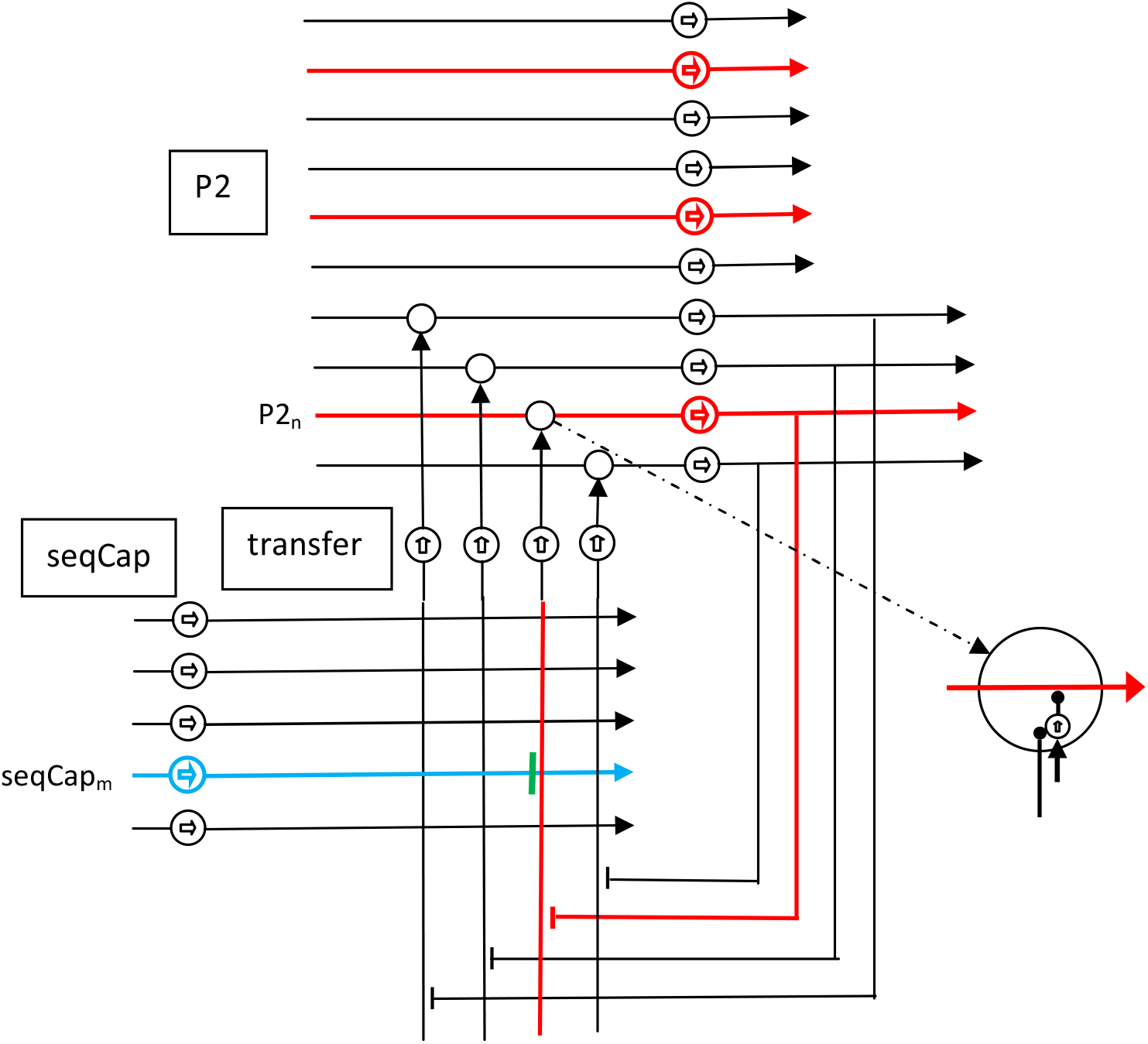
seqCap capture-only phase: red = activation pattern to be learned, blue = activation of cell (group) to learn pattern, green = seqCap_m_ synapse captures active P2_n_ output. An inhibitory version of a gating microcircuit shown in lower right. During the first forward pass the absence of excitatory input to the microcircuit interneuron suppresses inhibition to P2 dendrite, allowing distal excitation through ungated. The acquisition of P2 activation by one of the seqCap cells or cell groups is unaffected.

**Figure 10(b):**
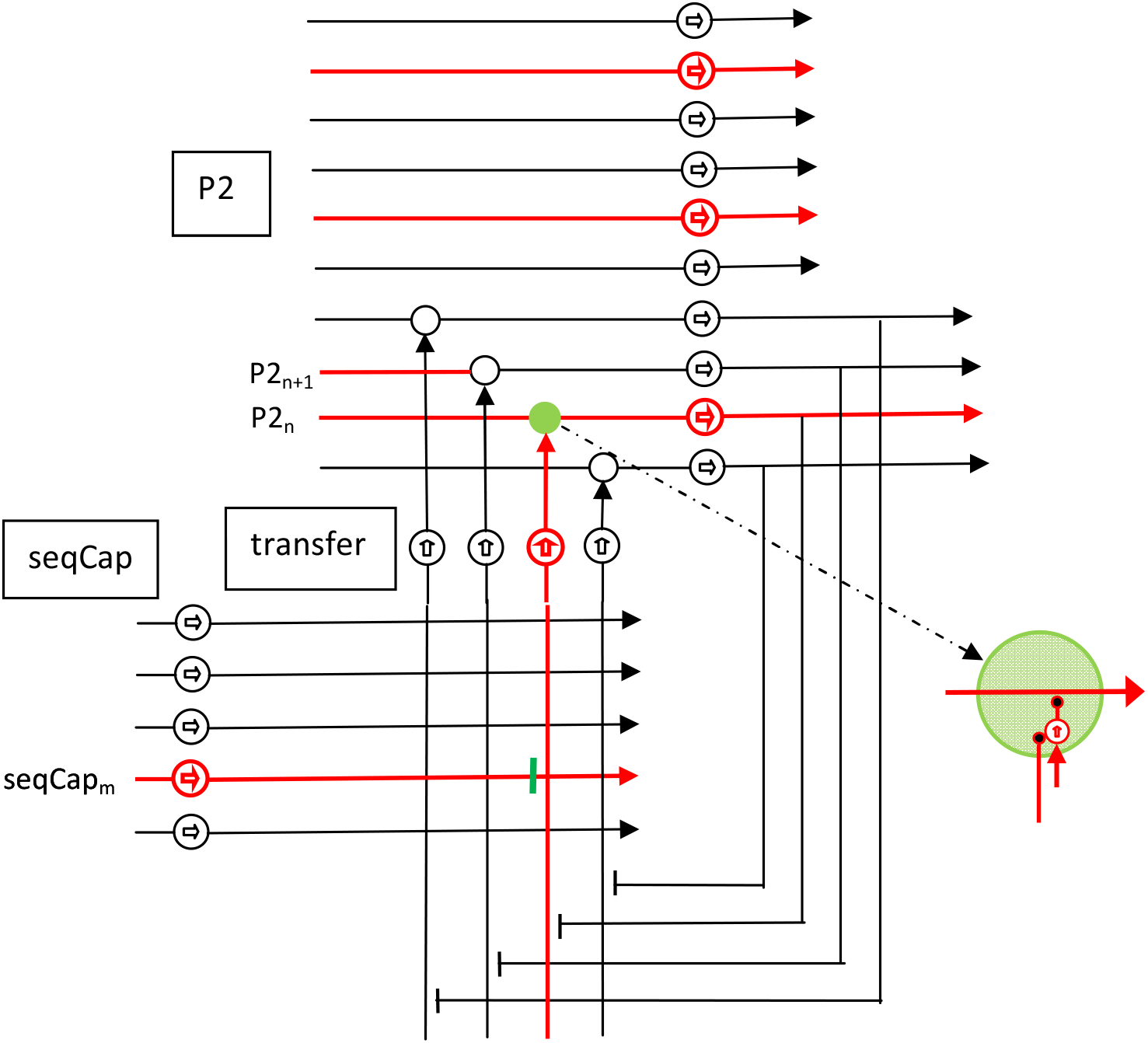
seqCap recall phase: red = seqCap_m_ cell and transfer cell activated, green = P2_n_ gating active. Inhibitory gating shown, in which active transfer axon inhibits “default” inhibition of P2 dendrite in the recall phase, thereby implementing a multiply or AND of the P1 mapping input to P2 and the seqCap recall activation. Mapping input to P2_n+1_ is active but blocked by the gating input from seqCap being inactive. Capture of the newly gated P2 state to seqCap follows recall of the previous P2 state from seqCap.

In Figure 9 the cell labeled “masking” represents an array of cells which provide an inhibitory pattern to block activation of P2 place cells in an excluded area. The functionality is that of biological boundary cells. (Boundary cells are found in subiculum, proximal to CA1. [7] This is downstream of where masking is indicated in Figure 9, but as their effect is applied on every cycle through the circuit the difference is negligible.) The initialization circuitry for this masking array is left undefined as the initialization is likely NOT provided through the main loop, but presumably directly via EC.

The remaining circuit feature that needs introduction is the goal-detected neuron. That cell has been discussed before in the simulation section. It is one neuron, or more likely a group of neurons whose dendrites span the axons of P2. Only the synapse from the axon of the P2 cell that is the goal of the path has non-zero weight. Hence, as discussed earlier, when the P2 activation reaches this cell, the goal-detected cell fires and triggers the events listed above that end the forward pass and initiate the backward pass. Among these is the switch of activation from the forward seqCap daisy-chain to the backward seqCap daisy-chain. It is most likely that the synapses of these neurons would be initialized by EC, and this would imply an input from EC to P2, which is not shown here but is a path often depicted in Hc-EC-complex connectivity diagrams.

#### Biological Hc Correlates

The dataflow in Figure 9 bears a strong resemblance to simplified renderings of Hc neuronal architecture. The general architectural similarity suggests the array of cells labeled P1 would be CA3 place cells and the array of cells labeled P2 would be CA1 place cells. The hypothesized connectivity between the P1 axons and P2 dendrites implementing the mappings described in both navigational and relational modes constitute a more specific architecture than is known of the actual connection between CA3 and CA1. Many renderings of Hc circuitry depict multiple inputs to CA1 beside CA3. There is no current evidence to confirm or refute the assumption here that among those inputs are memory structures for the capture of transient activity of CA3 or CA1, here labeled SequenceCapture. But that such a structure exists within the Hc complex would be hard to argue against given the experimental evidence for bidirectional playback of place cell sequences comprising experienced journeys. [3] The circuit elements that capture those sequences would almost certainly have to capture them from the dendrites of CA1 (or P2 as labeled here). The goal-detection function on the outputs of P2 is consistent with the experimental identification of CA1 as the locus of navigation goal targets. [9]

While the diagrams in Figure 1 and Figure 9 show the outputs of P2/CA1 as direct inputs to P1/CA3, this is apparently an over-simplification. Most anatomical depictions of the Hc-EC complex do not render direct connections within Hc that would implement that path. Instead most depict the outputs of CA1 feeding subiculum, EC and several other structures, which then feed CA3. So the implication is that the loop is closed outside of Hc proper. However, some evidence of a direct connection does exist in the form of an experiment in which EC is disabled but Hc signals continue to loop.

Note that the recurrent connections within CA3 play no role in this hypothesis.

Since no biological structure can be associated confidently with the sequence capture architecture hypothesized here, and there are multiple possible implementations to achieve the functionality described in the simulation section, one can only hedge on its neuronal implementation. SeqCap inputs to P2 dendrites must act as gating, such that those connections which correspond to non-zero states of SeqCap cell axon synapses allow signals from the mapping connections from P1 to reach the somas of P2, and the zero states block the mapped signals from P1. This behavior can be implemented most simply by “inverting” the states of the SeqCap axon synapses and using the inverted states to induce shunting inhibition in P2 dendrites, or somatic inhibition. The microcircuitry to accomplish this is left ambiguous in Figure 9, but there are several alternatives, one of which is presented in more detail in Appendix 3. And it is possible, though maybe less likely, that the gating could be implemented by multiplicative excitatory inputs at critical locations on the dendrites.

The simulations assumed, for clarity, an array of SeqCap cells as large as P2 for each path step to be captured and replayed for intersection. There are alternative architectures which would be functionally equivalent but require far fewer cells by storing the memorized states in the synapses between seqCap cells and some local microcircuitry leading to Inhibitory inputs to P2 place cells. A simple version of a possible architecture is presented next.

#### Dense seqCap memory and “inverting” microcircuitry

Instead of implementing each seqCap as a array of the same dimension as P2 (as described for the simulation), seqCap can be implemented more densely by holding the P2 activation value in synapses between one (or a few) seqCap cell(s) and a set of transfer interneurons, each of which drives the gating synapse on one P2 dendrite. Figures 10(a, b) illustrate a detail of the circuit in Figure 9 using a single seqCap cell for each P2 capture and recall. More realistically, each illustrated seqCap cell represents several, each of whose axons synapse on a subset of the transfer cells. This keeps synapse counts realistic while providing redundancy.

During the first forward pass there is no intersection operation because seqCap hasn’t received any activation patterns from P2. Therefore the mapped P1 activations must be passed without gating to the P2 soma. But during this pass seqCap must also receive the outputs from P2. This is in effect a capture-only phase and is illustrated in Figure 10(a). The caption explains the operation of the microcircuit. The daisy-chain selection of seqCap cell or cell groups is not illustrated, but is the same as in Figure 9.

Starting with the first backward pass, all passes incorporate the intersection operation between the mapped P1 excitations of the P2 dendrites and the activation pattern recalled from one of the seqCap cells or cell groups. During these passes a continuous excitatory input to the inhibitory interneuron of he microcircuit enables shunting inhibition to the target P2 dendrite **only if** there is no inhibitory input to the interneuron from the axons of the selected seqCap cell or cells. Hence active seqCap axons via the transfer interneurons inhibit the inhibitory interneuron, allowing the distal activation of the P2 dendrite to reach the soma, as indicated by the color coding in Figure 10(b). This implements the desired gating during all passes after the first forward pass. The capture of P2 output by seqCap replaces the pattern stored during the previous pass in the seqCap synapses. The timing of the “read” from seqCap and the “write to” seqCap is controlled by the timing of the excitation on the selected seqCap dendrite and must insure the recall/intersection operation is complete before the capture by seqCap takes place and changes the synapse weights. Whether the update of the selected seqCap synapses with the transfer interneurons by short-term facilitation and depression can take place fast enough is not determinable from current available experimental results.

#### Alternative short-term memory architectures for seqCap

The viability of the dense seqCap memory architecture of Figure 10(a,b) is dependent on the time scale of the fastest short-term potentiation (STP) processes. The required persistence of the memory state can vary significantly depending on the number of steps in the pass, from a few to dozens. Observed STP processes appear to have sufficient duration. [10]

Several alternative architectures are possible for the dense architecture which relax some of the timing requirements using extra neurons in the path as buffers in effect. These would still functionally remain in the category of the architecture of Figure 10(a,b).

A less critical alternative architecture keeps the large seqCap cell array of the simulation architecture.

In this the captured pattern resides in the oscillatory state of each cell in the array. Each seqCap array is gated onto the transfer interneurons as it is selected by the daisy-chain described earlier. In this arrangement, like that of Figure 10(a,b) the state of the seqCap cells has to be inverted to control shunting inhibition on the P2 dendrites. Hence the inhibitory interneuron microcircuitry seen in Figure 10(a,b) and its function would remain exactly the same.

Figure 11 illustrates a possible architecture for the cell array architecture for seqCap and its connections to P2. The seqCap_m_ cells shown are four of several thousand (for the simulations above), one per place cell in P2. Before each capture they are silenced by general inhibition, then a subset are set in an oscillatory state by the active inputs from P2 axons. The selection input from the associated daisy-chain cell determines which seqCap array is affected. On recall/intersection an excitatory input to an interneuron microcircuit gates the seqCap_m_ array onto the transfer neurons. The overall functionality from the perspective of the P1-P2-P1 loop is identical to the circuit detail of Figure 10(a,b).

**Figure 11:**
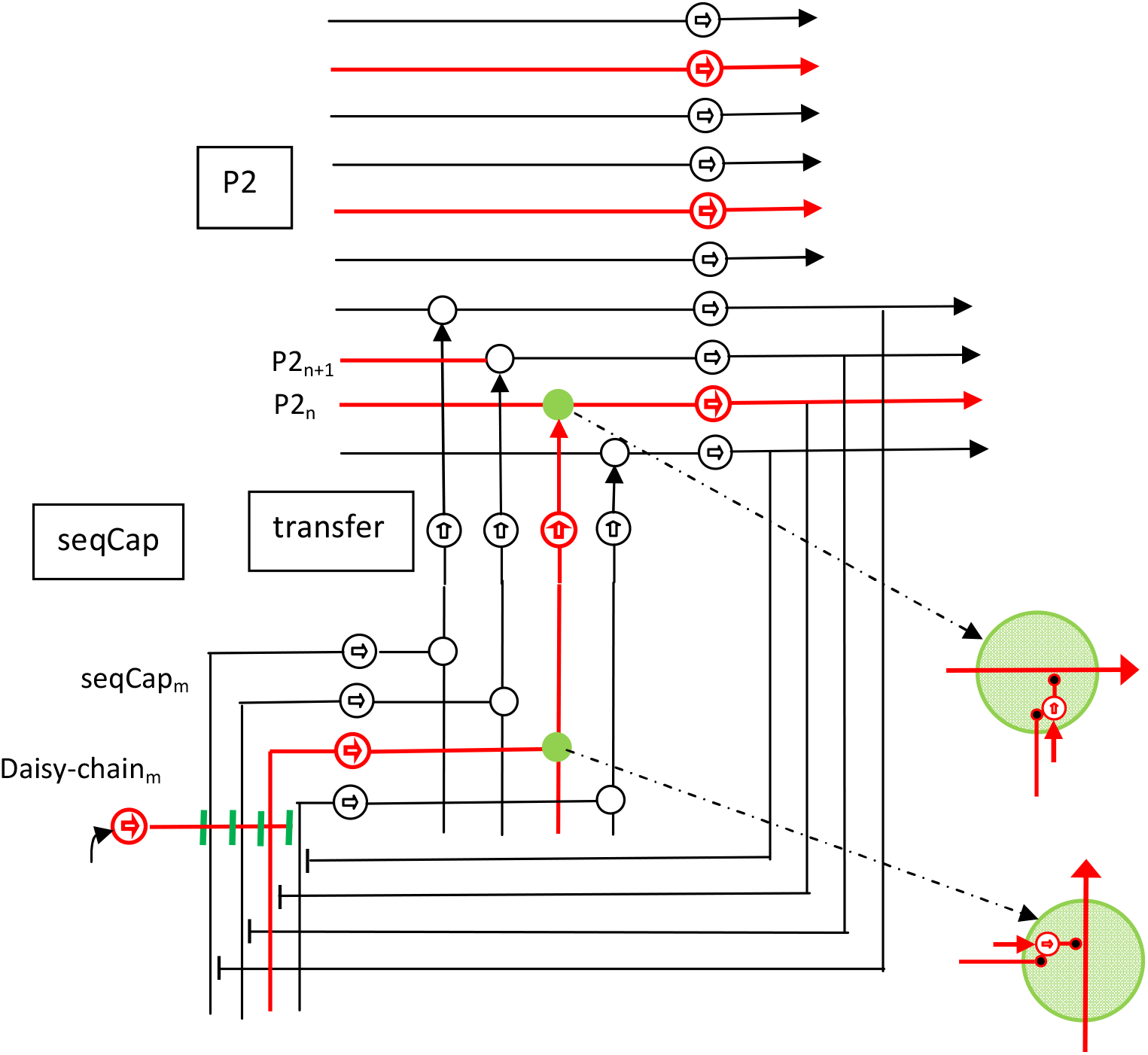
seqCap circuit detail for a cell array architecture. Recall/intersection activation shown in red.

There is no evidence this author is aware of at present to further resolve which architecture is most likely to be involved in biological Hc.

#### Boundary or Border Cells

Boundary or border cells, as reported in the experimental literature, fire when the animal is within a certain distance of a boundary. [7] In the planning process, the interaction must somewhat different. In order to influence a route which the animal has not yet taken. The presence of a boundary or barrier must suppress the place cells near, at, and beyond the border. In the context of path planning this implies boundary cells would be caused to fire when the place cells of the hypothesized route come into range. But then their firing must act in such a way as influence the convergence of the path-planning by blocking the path solution from crossing the border cell defined regions. This can be accomplished either by direct inhibition of the place cells in the region of the boundary cell or, indirectly, by activating co-located cells in an array (like the mask array defining the obstacles for Figures 5 and 6) which in turn inhibit activation of place cells.

The navigation simulations above utilized a mask array to block out whole excluded areas, though, as noted in the earlier discussion, the mask array could just as well have been initialized to exclude just the border of the prohibited area, so long as that border zone was wider than the longest mapping. Animals, however, are unlikely to be aware of excluded regions as seen from above, but rather observe only barrier surfaces. The literature indicates that border cells represent the spatial locations of such barriers. Merely masking the P2 place cells at the location of the barrier is not sufficient as the mappings that jump more than one cell would usually not be blocked. Consequently, a single boundary cell’s masking influence would have to inhibit a radius in CA1/P2 greater than the longest mapping. For this reason the details of the mask cells’ connectivity is not rendered in Figure 9.

#### Relationship Circuit

The extension of the navigation circuit to compute the chained inferences discussed earlier requires changing one step in the algorithm: the replacement of the static local radial displacement (translation) mappings with a mechanism for retrieving the relevant pairs of source-destination place cell loci that implement the allowed mappings. From a circuit perspective this involves detecting the source locus for each relevant pair on the axons of P1 and inducing the destination loci on the dendrites of P2 for each pair whose source locus corresponds to an active axon of P1. In Figure 12, this mechanism is depicted as implemented as two coupled cells: one to detect the source locus on P1 and one to induce the destination locus on P2. The mappings for the route planning mode are either disabled (which would require circuitry similar to that for path integration, as above), or the mapping for the relational computation takes place on a different set of Hc place cells. It is also possible for the two mapping mechanisms to co-exist in the same space so long as the chain of inferences is quite short and the space between assignments of place cells to entities is much longer than the longest distance traversed in a few steps by the route planning mappings.

**Figure 12:**
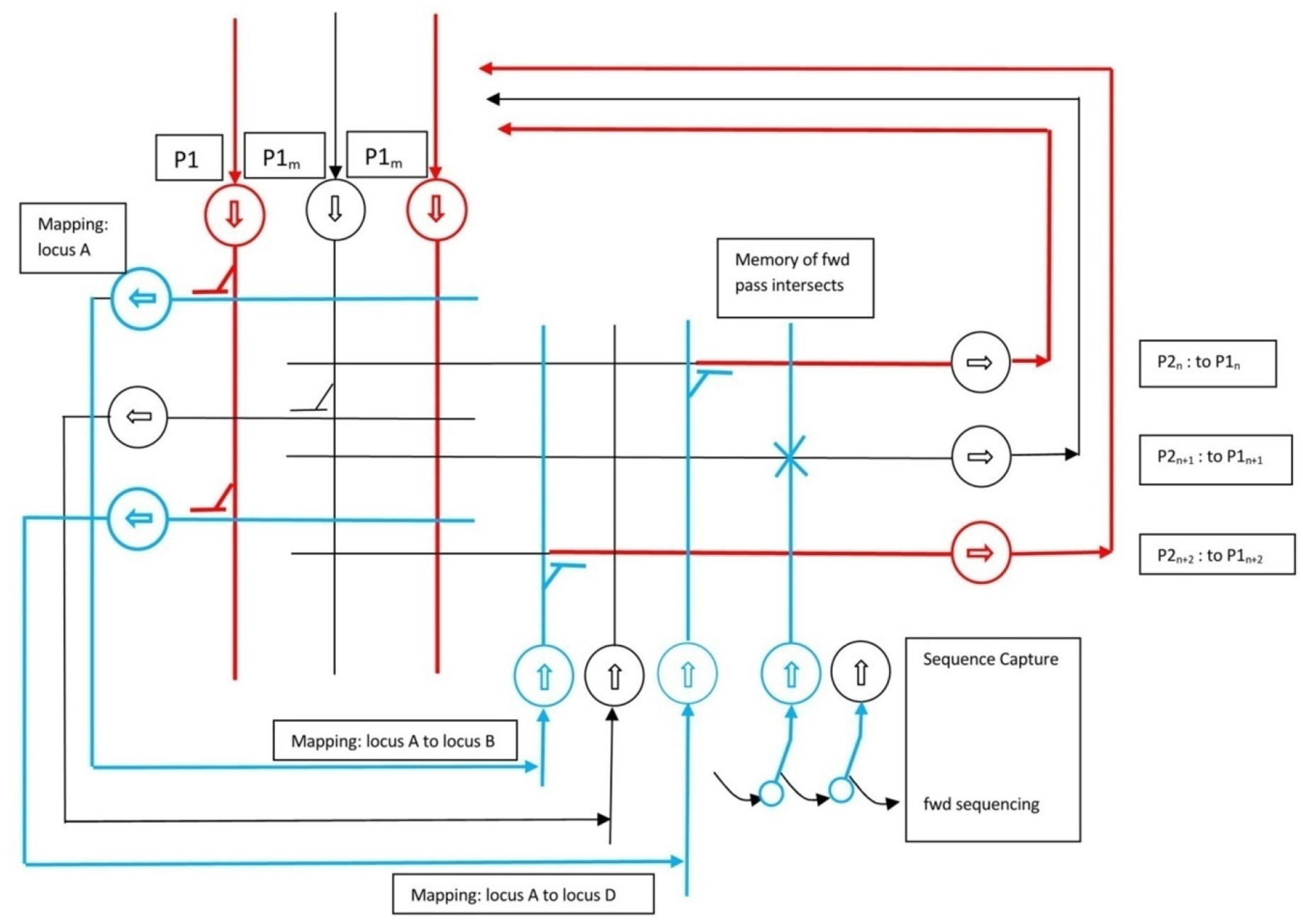
Relational mapping mechanism on the same circuit as in Figure 9. For simplicity only the forward sequencing daisy chain is shown, but both directions are required, as in shown in Figure 9, and discussed in the relational process.

In Figure 12, aside from the cell pairs to implement the relationship mappings, the circuit is identical to the navigation circuit. In particular seqCap plays the same role as in navigation. For the last backward pass step the intersection of P2 by seqCap is suppressed by the absence of a seqCap array in the selection daisy chain for that sequence step, as discussed in the section on the navigational circuit.

## Discussion

### Plausibility of the hypothesis

How plausible is the foregoing hypothesis? There are two critical elements which make the simple P1-P2-P1 loop capable of the solutions illustrated. They are the mapping connectivity from P1 to P2 and the P2 activation sequence memory seqCap.

The support for something like the seqCap structure is strong but partial. That there exists a component of the Hc circuitry that can save the loci of a path sequence and play it back in either direction has been well demonstrated. But the hypothesized role of providing the pruning by intersection P2 activity to the path or relational chain solution is not supported by any neuroanatomical evidence of which the author is aware. Nor is the proposal refuted by any evidence of which the author is aware. The functional argument in support, that there do not yet appear to be functional alternatives, still leaves it a pure hypothesis. At least it gives the experimentalist something specific to look for, if adequate techniques exist or when they do.

The second critical element is the mapping connectivity. This would be best confirmed by direct anatomical evidence, but that of course is out of reach at this time. The second best evidence would be activity recordings of sufficient resolution to confirm the convergence dynamics. The panels of Figure 2 impart a sense of what neural recordings of CA3 and CA1 might look like in an experimental context in which a path planning problem was posed. Experimental confirmation would be difficult given the current limitation of number of recording channels on the one hand and the temporal resolution limitations of imaging on the other. To record the initial spread of activation is practical, but recording the converged state would require having electrode in exactly the right neurons for all or much of the converges path. Recordings in some experimental contexts show burst activity consistent with the initial spread of activation and periodic excitation consistent with multistep dynamics in the simulation.[11] Similarly, evidence of the limited span of excitation in CA1 from individual CA3 cells [11] is consistent with but not conclusive of the mapping proposed in this paper. So it can at most be considered suggestive.

### Relevance of the hypothesized function to other Hc roles

The relevance of the proposed mechanism to the extensive evidence of Hc’s role in memory has not been addressed. The only recall function included is the sequenceCapture structure, necessary to the convergence dynamics. But this is an exceedingly specialized memory function, and not in any obvious way related to the variety of memory roles played by Hc.

If there is an Hc functionality explored above that might have relevance to the various memory roles played by Hc, it is the relational mechanism in the latter half of the paper. The ability to test novel relational conjectures by the discovery of inference chains is applicable well beyond the transitive preference experiments which confirmed the capability and Hc’s role in it. Causal relationships, temporal relationships, hierarchical relationships, for example, are all representable by directed graphs, which are a generalization of what the mechanism presented above explores. Memories, to be useful, need context, and relationships are the organization of context. [12] So while nothing beyond the simple abstract ordering problem is presented above, the implications of the mechanism’s capability appear to be much broader.

### Relationship to Map-Seeking Circuit (MSC)

Readers familiar with the author’s earlier work on the MSC algorithm and circuit [13] may have noticed the common element of forward and backward dataflow interaction. But this is where the commonality ends. In MSC that interaction is comprised of measurements of similarity between the flows at different stages which then drive coefficients governing the contribution of different mappings. There is no direct interaction of the flows by an intersection operation which is the crucial element of the circuits described in this paper. While MSC has been shown to be capable of path planning computations [14], MSC as always described has a fixed number of layers, which is not suitable for solution of paths of unknown length. The MSC computation can be implemented similarly to the circuit and algorithm described here, as a loop with storage like seqCap, and has been given the working moniker rMSC for “re-entrant MSC.” But the rMSC dynamic still requires the measurements between forward and backward flows and coefficients to control the contribution of each mapping at each step of the computation. The circuit and algorithm described here is much less compute/circuitry intensive than a looping MSC configured for the same purpose and converges faster in most conditions.

## Appendix 1: PseudoCode

The algorithmic steps for one iteration of path planning is shown below. For multiple iterations the following code is repeated with the contents of seqCap from the last pass of one iteration carried over to the first pass of the next iteration.

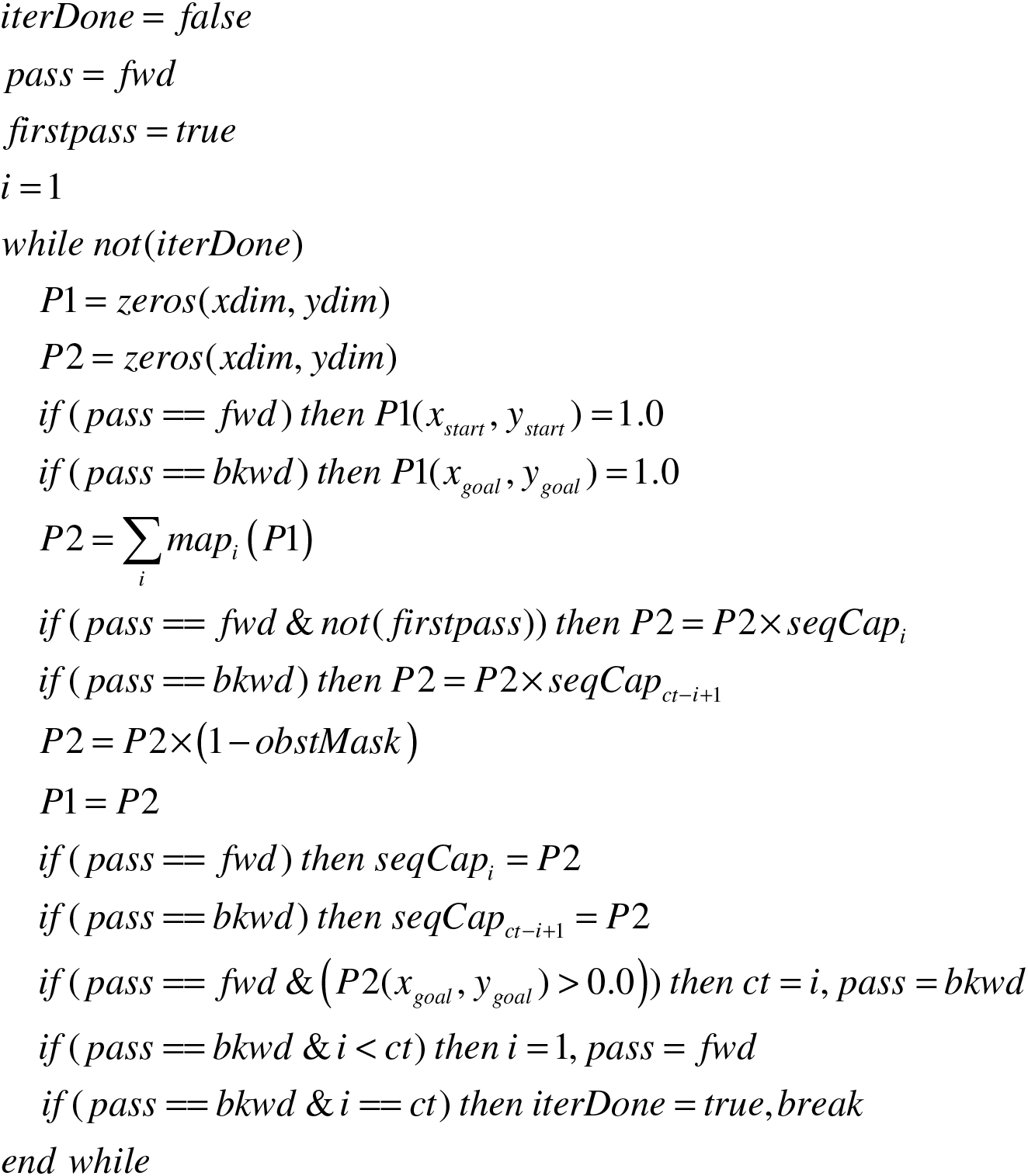

### The algorithmic steps for computing the causal chain for multi-degree ordered relationships is shown below. (Only one iteration is required.)

The mapping operation *map_i_*() below implements the P1 axon to P2 dendrite connectivity that establishes ordered relationships, e.g. A *rel* B or *relation(A,* B). Each relation A *rel* B is represented by a pair [A B]. A and B are place cell arrays conforming to P1 and P2 in which the the place cell representing entity A = 1.0 and the rest of A= 0.0, and the place cell representing entity B = 1.0 and the rest of B = 0.0. So for each *map_i_* if there is a non-zero place cell in P1 corresponding to the non-zero place cell of A, then the place cell in P2 corresponding to the non-zero place cell of B has 1.0 added to it.

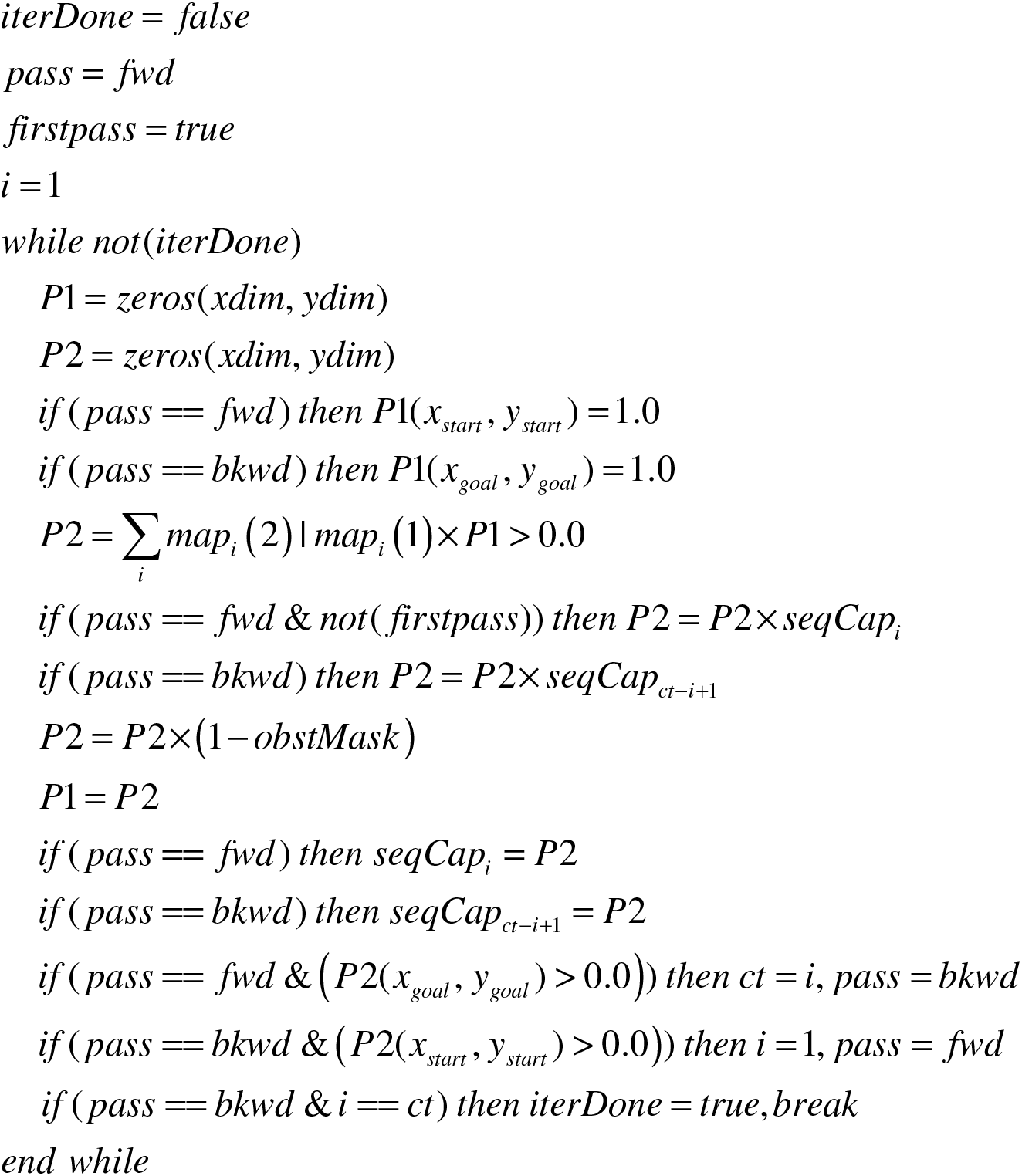

## Appendix 2 Path Integration

The process described above solves the inverse problem of finding an efficient route between a starting location and a known goal location under the constraints of obstacles or available pathways. There is also evidence, perhaps not conclusive, that Hc is also involved in the solution of the forward problem associated with the inverse problem of route finding, sometimes referred to as *path integration* or *odometry.* This is the incremental computing of the likely present position based on the speed and direction of small intervals of the journey. The architecture presented above requires a minor modification to be capable of that computation. Instead of being always engaged the mappings from P1 to P2 need to be selectable as to direction and length based on known heading and speed of travel. Algorithmically this is a trivial modification. Modification to the neuronal circuit involves introducing a control neuron for each mapping angle and distance combination which gates a selected mapping from P1 to P2. One control neuron controls all the mappings of a particular angle and distance from all place cells in P1, so the path is constructed by a sequence of selections as the path advances, driven by orientation and speed inputs from elsewhere in the brain. The neuronal circuit detail to implement this will be discussed briefly in a later section.

Effective path integration requires accurate specification of direction and speed, so a great number of mappings with fine discretization must be implemented from each location in P1 to a location in P2. As demonstrated above, the inverse problem of route finding does not require fine discretization since mappings which take a path diverging from the optimal direction for one or more steps, will automatically be compensated by the seeking process which minimizes the number of steps, given the mappings available. Since the primary focus of this paper are the inverse problems solved by Hc, discussions of the complexities of path integration, such as error correction, will not be discussed.

### Path Integration Circuit Modifications

As discussed earlier, if the same part of the Hc circuit that computes the inverse problem of route planning is also engaged in the forward problem of path integration, then the mapping circuitry has to be modified to allow specific maps to be enabled at each step of the journey. Therefore, instead of mapping connections which are always enabled, the mappings for particular angle and distance combinations must be selectable. Figure X shows a possible circuit arrangement to enable this. Each mapping connection from P1 to P2 that executes a particular angle-distance combination only becomes enabled if the mapping cell associated with that transformation is enabled. These are labeled map-selection neurons. They provide activation to a synapse paired with the mapping synapse between P1 and P2, or some equivalent circuitry. Both synapses of the pair, or equivalent, must receive activation for the mapping to be effected. This is another multiplicative or gating behavior. A single map-selection neuron can provide the control for a large number of common angle-distance mappings between P1 and P2. (Note.This a simplified version of the dynamic mapping of the map-seeking-circuit whose operation has been demonstrated in analog circuitry. [])

As discussed earlier, path integration requires much more finely discretized angle-distance mappings than route planning, and the requisite fine grain spatial representation. So it is possible that this forward problem is solved in a different area, or on a different resolution place cell set, of Hc, and the selectable mappings coexist with the fixed mappings that are sufficient for the inverse route planning problem.

**Figure A2-1:**
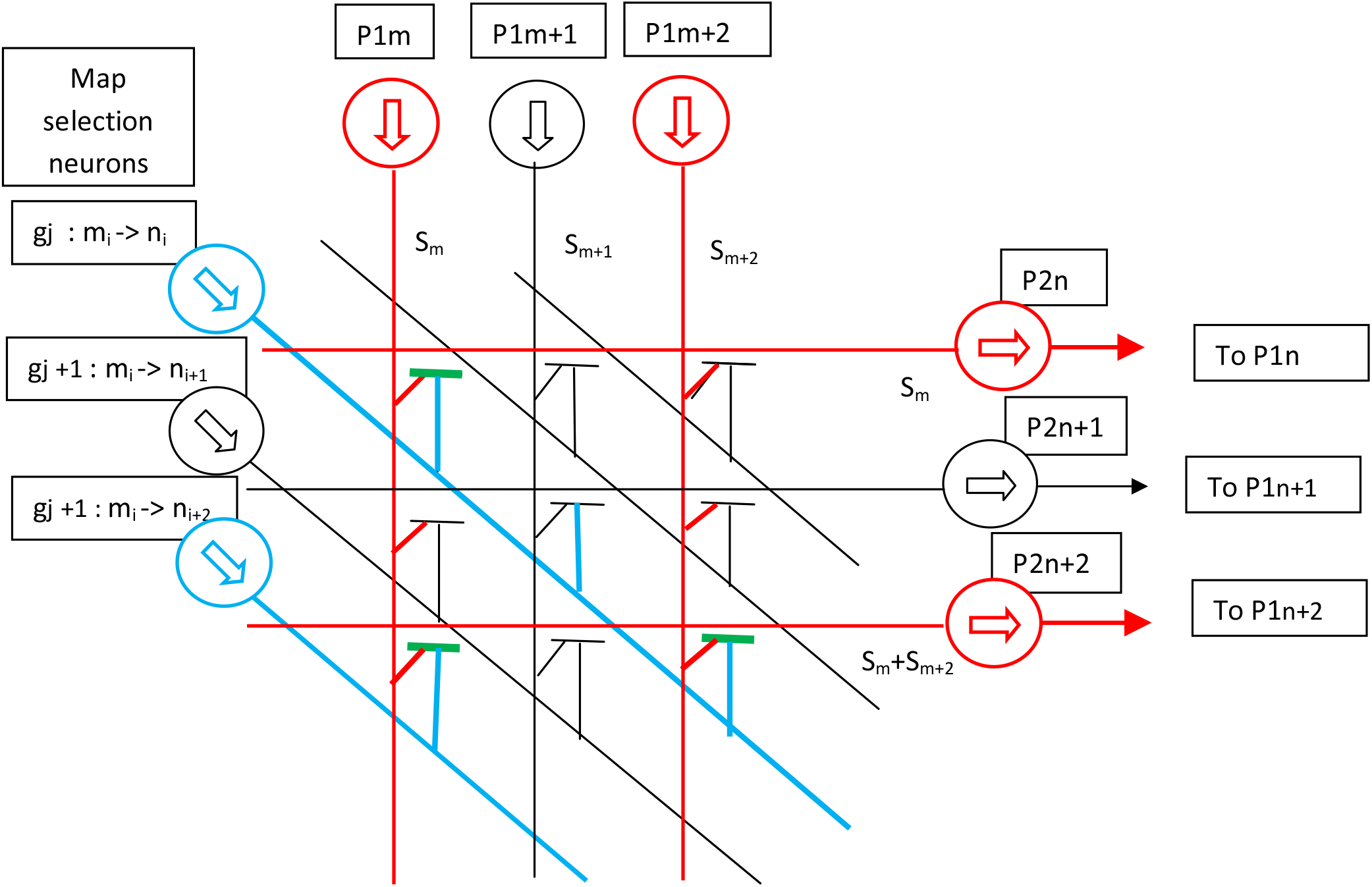
Selectable P1 to P2 mappings for path integration

